# BET Bromodomain Proteins Regulate Transcriptional Reprogramming in Genetic Dilated Cardiomyopathy

**DOI:** 10.1101/2020.02.09.940882

**Authors:** Andrew Antolic, Hiroko Wakimoto, Zhe Jiao, Joshua M. Gorham, Steven R. DePalma, David A. Conner, Da Young Lee, Jun Qi, Jonathan G. Seidman, James E. Bradner, Jonathan D. Brown, Saptarsi M. Haldar, Christine E. Seidman, Michael A. Burke

**Affiliations:** Emory University School of Medicine; Harvard Medical School; Dana Farber Cancer Institute; Brigham and Women’s Hospital; Vanderbilt School of Medicine; Gladstone Institute of Cardiovascular Disease; Department of Medicine, Cardiology Division, UCSF School of Medicine; Amgen Research; Howard Hughes Medical Institute

## Abstract

The bromodomain and extraterminal (BET) family of epigenetic reader proteins are key regulators of pathologic gene expression in the heart. Using mice carrying a human mutation in phospholamban (PLN^R9C^) that develop progressive dilated cardiomyopathy (DCM), we previously identified the activation of inflammatory gene networks as a key early driver of DCM. We reasoned that BETs control this inflammatory process, representing a key node in the progression of genetic DCM. Using a chemical genetic strategy, PLN^R9C^ or age-matched wild type mice were treated longitudinally with the BET inhibitor JQ1 or vehicle. JQ1 abrogated DCM, reduced cardiac fibrosis, and prolonged survival in PLN^R9C^ mice by inhibiting inflammatory gene network expression at all disease stages. Cardiac fibroblast proliferation was also substantially reduced by JQ1. Interestingly, JQ1 had profound effects on pathologic gene network expression in cardiac fibroblasts, while having little effect on transcription in cardiomyocytes. Using co-immunoprecipitation, we identified BRD4 as a direct and essential regulator of NFκB-mediated inflammatory gene transcription in cardiac fibroblasts. In this this model of chronic, heritable DCM, BETs activate inflammatory gene networks in cardiac fibroblasts via an NFκB-dependent mechanism, marking them as critical effectors of pathologic gene expression.

## INTRODUCTION

Heart failure (HF) is a common and morbid disease. Existing pharmacotherapies have improved clinical outcomes for HF patients, but residual mortality remains exceedingly high with an estimated 5-year survival of approximately 50%.(1) HF often results from dilated cardiomyopathy (DCM), a heterogeneous disease with a strong genetic basis.(2) Characteristic phenotypic features of DCM include ventricular wall thinning, cardiac chamber dilatation, eccentric myocyte hypertrophy, diffuse interstitial fibrosis and reduced systolic function. Concomitant with this negative cardiac remodeling is the activation of a suite of cardiac signaling cascades that result in pathologic gene expression. These dynamic changes in gene expression are coordinately controlled in temporal fashion at different stages of disease.(3)

To study the regulation of pathologic gene expression in DCM, we utilized a transgenic mouse model expressing a missense mutation (p.Arg9Cys) in phospholamban (PLN^R9C^), a dynamically regulated protein that controls calcium cycling via regulation of the cardiac sarcoplasmic/endoplasmic reticulum calcium adenosine triphosphate (SERCA2a) pump.(4) Abnormal cardiomyocyte calcium homeostasis is a common and proximal mediator of stress-induced cardiac remodeling.(5, 6) PLN^R9C^ mice display altered calcium handling prior to onset of phenotypic disease (preDCM) and ultimately develop progressive DCM with fulminant heart failure culminating in premature death.(4) This inexorable process mimics the disease course of individuals carrying this mutation(4, 7, 8) and recapitulates the natural history of DCM. Thus, PLN^R9C^ is a powerful model by which to study the molecular changes that underlie DCM.

We previously demonstrated temporal changes in gene expression in PLN^R9C^ hearts.(3) These findings were similar to those seen in human induced pluripotent stem cells engineered to carry the PLN^R9C^ mutation,(9) and highlight the coordinate integration of multiple signaling networks into a concentrated and temporal stress-response program. Pathologic gene networks activated in HF are associated with dynamic remodeling of chromatin(10, 11), including changes in lysine acetylation of histone amino-terminal tails and other chromatin-associated proteins. These acetylated lysine residues can be recognized by epigenetic reader proteins harboring acetyl-lysine recognition domains (bromodomains), which in turn regulate gene transcription. The bromodomain and extraterminal (BET) family of epigenetic reader proteins (BRD2, BRD3 and BRD4) bind to acetylated chromatin via their dual N-terminal bromodomains and co-activate gene transcription by assembling protein complexes that promote pause release of RNA-polymerase.(12, 13) BETs have been identified as critical coactivators of pathologic gene expression in rodent models of pressure-overload and ischemia-mediated heart failure (13–15). Gene expression profiling in these models suggest that BETs exert therapeutic effects via preferential suppression of inflammatory and pro-fibrotic transcriptional programs.

The PLN^R9C^ mouse model features robust activation of a broad array of inflammatory gene networks very early in the disease course, prior to the onset of LV dilation or systolic dysfunction.(3) However, it remains unknown whether activation of this transcriptional program plays a causal role in DCM pathogenesis. Interestingly, the genes and networks upregulated in PLN^R9C^ mice were remarkably similar to those suppressed by JQ1 treatment in other mouse cardiomyopathy models. Therefore, we tested the hypothesis that inhibition of BETs with the small molecule JQ1 could directly suppress pathologic inflammatory gene expression and thereby exert salutary effects in this model of chronic, genetic DCM. Herein, we demonstrate the effectiveness of JQ1 in suppressing inflammatory gene network activation, thus establishing a causal relationship between early pathologic gene expression and DCM progression. We identify cardiac fibroblasts as the chief driver of this process, and substantiate the critical mechanistic link between BRD4 and NFκB, establishing BETs as nodal activators of cardiac fibroblasts in DCM.

## METHODS

### Mouse model and drug treatment

Mice were housed in ventilated racks in a temperature and humidity controlled, pathogen-free facility with 12-hour light-dark cycles and free access to water and standard laboratory mouse chow. Transgenic mice overexpressing PLN^R9C^ under control of the α-cardiac myosin heavy chain promoter in the FVB genetic background have been characterized.(4) Male PLN^R9C^ mice were compared to age-matched wild type (WT) male FVB control mice.

The primary endpoint of mouse survival was defined as survival free of overt HF. PLN^R9C^ mice ultimately develop HF (increased respiratory rate, decreased activity and ability to withdraw from touch, cachexia and cool skin) in the final stages of disease. Mice die within 3-5 days of symptom onset, marking these symptoms as a surrogate for impending death such that mice were sacrificed to prevent unnecessary pain or suffering.

Stock JQ1 was prepared as previously described.(13) Mice were treated with JQ1 (50mg/kg) or vehicle (10% DMSO in 10% hydroxypropyl β-cyclodextrin) daily via intraperitoneal injection. The length of dose administration varied depending on the study question. Hearts were harvested under deep anesthesia and LV tissue was extracted for experiments as detailed in the **Supplementary Methods**.

### Study approval

All studies were approved by the Institutional Animal Care and Use Committees of Harvard Medical School and Emory University, and performed in accordance with the NIH Guide for Care and Use of Laboratory Animals.

### Echocardiography

Mice were anesthetized using an isoflurane vaporizer and attached to ECG leads on a Vevo Mouse Handling Table. Chest hair was removed with depilatory cream (Nair). Transthoracic echocardiography was performed with heart rate >500bpm using a Vevo 770 High-Resolution In Vivo Micro-Imaging System and RMV 707B scan-head. A single, experienced echocardiographer (HW) blinded to genotype, acquired the images and analyzed the data. Parasternal 2D images and M-mode images were acquired to assess (1) LV end diastolic diameter (LVEDD), (2) LV end systolic diameter (LVESD), (3) LV wall thickness (LVWT; the combined anterior and posterior wall thickness), and (4) fractional shortening (calculated as FS=(LVEDD -LVESD)/LVEDD)).

### Histochemistry

Ventricular tissue was fixed in 4% paraformaldehyde, paraffin-embedded and sections cut from apex to base to cover all regions of the LV (see **Supplementary Methods** for additional details on all histochemical protocols). Nuclei were stained with DAPI (Sigma, D9542), and the extracellular matrix was stained with anti-wheat germ agglutinin (WGA; Molecular Probes, W32464). To quantify fibrosis, sections were stained with Masson trichrome and the ratio of fibrotic area to total ventricular area was measured using a Keyence BZ-X700 microscope.

To assess cell proliferation mice were injected intraperitoneally on 2 consecutive days prior to sacrifice with 5-ethynyl-2’-deoxyuracil (EdU) at 25mg/kg body weight in 10% DMSO-PBS at 14-weeks of age (**Supplementary Figure 1**). Tissue was stained using the Click-iT assay (Thermo-Fisher) and confocal microscopy performed to quantify EdU-positive cells. To identify the non-myocyte cells that were EdU-positive, additional slides were co-stained with the Click-iT assay and either anti-vimentin (Abcam, ab92547), anti-CD31 (Abcam, ab124432) or anti-CD45 (Abcam, ab10558).

### Cell Isolation

Myocyte and non-myocyte cell populations were isolated from PLN^R9C/+^ and age-matched wild type mice treated with either JQ1 or vehicle at 16-weeks of age (n=3/treatment group). Hearts were excised and retrograde coronary perfusion was established via aortic cannulation.(16) The heart was perfused with enzyme buffer (see **Supplementary Methods**) for 10 minutes. The atria and right ventricle were removed and the LV was minced into small pieces in transfer buffer (perfusion solution + 5mg/mL BSA) and then passed several times through a sterile pipette. The resulting cell suspension was passed through a mesh filter into a 50mL centrifuge tube and incubated for 15 minutes at RT to allow myocytes to pellet by gravity. The pellet was collected as a cell fraction enriched in myocytes. The supernatant from the filtered cell solution was centrifuged at 1500rpm for 5 minutes at 4°C, and then plated in 75mm tissue culture dishes. After 2 hours, dead cells were washed off with 1x PBS and plated cells collected, resulting in a non-myocyte cell population that is enriched for cardiac fibroblasts (**Supplementary Figure 2**).

### RNA-seq

RNA was extracted from LV tissue or pooled cardiac fibroblast or cardiomyocyte fractions using TRIzol. Prior to cDNA synthesis, 2 rounds of poly-A selection were performed using oligo dT Dynabeads (Invitrogen). cDNA libraries were constructed from individual mouse samples and RNA-seq libraries were constructed using the Nextera XT DNA Library Preparation Kit (Illumina). Libraries were sequenced on the Illumina platform, then aligned to the mouse reference sequence mm10 using the STAR aligner software(17) and a custom data processing pipeline as previously described.(3)

### Bioinformatics Pipeline for Pathway Analysis

Genes were defined as differentially expressed if they met ALL of the following 3 criteria: (1) normalized read count >1 copy/million reads in the RNA-seq library, and (2) fold-change >1.33 (up-regulated) or < -1.33 (down-regulated), and (3) p-value <0.001 for the comparison of normalized read counts between experimental samples of interest. To increase stringency, all genes identified as differentially expressed had to meet all of these parameters for all comparisons between n=3 mice in each group (total of 9 comparisons per gene). Differentially expressed genes meeting these parameters were identified using a custom pipeline in R (version 3.0.1). Culled lists of differentially expressed genes were then subjected to bioinformatics analyses using Ingenuity Pathway Analysis (IPA) as described in detail in the **Supplementary Methods**.

### Western blotting and immunoprecipitation

Total protein was extracted using RIPA buffer, quantified, separated by SDS-polyacrylamide gel electrophoresis (PAGE) on 4-12% precast bis-tris gels and transferred to PVDF membranes. Western blotting was performed using antibodies to BRD2, BRD3, BRD4, vimentin, RELA and acetyl lysine-310 (aK310) RELA as described in the **Supplementary Methods**.

Immunoprecipitation (IP) was performed targeting BRD4 and the NFκB subunit p65/RELA. Freshly harvested whole left ventricles were dissolved in lysis buffer (50mM Tris-HCl, 250mM NaCl, 1% NP-40, 1mM EDTA) and 1mg of protein lysate was subjected to IP with 5ug of anti-BRD4 antibody (Abcam, #ab128874) or anti-NFκB p65 antibody (Abcam, #ab16502) overnight at 4°C with rotation.

Protein G magnetic beads (50μl) were added to the protein slurry and incubated for 3 hours with rotation. Beads were then washed twice in lysis buffer followed by elution in Laemmli buffer. SDS-PAGE and immunoblotting was performed as described above and using 1:500 anti-BRD4, 1:1000 anti-NFκB p65, or 1:1000 anti-NFκB p65 acetyl K310 (Abcam, #ab19870); 1:500 VeriBlot (Abcam, #ab131366) was used as the secondary antibody for all IP detection experiments.

### Statistics

Data are presented as mean ± standard deviation (SD). For echocardiography data, time-dependent between-groups variance was calculated using ANOVA with a model-based fixed-effects standard error method. To estimate sample size, a power calculation was performed using the “pwr” package in R based on our prior echo data in PLN^R9C^ mice.(3) To have 80% power to detect a 50% improvement in FS and LVEDD (a moderate effect size) with 4 groups of mice at a significance level of 0.05, we needed n=12 mice/group. Since PLN^R9C^ mice can die suddenly at any point, we used n=14 PLN^R9C^ mice in each arm. As JQ1 was not expected to alter echocardiographic parameters in WT mice, we reduced these arms to n=7. Tests of the 4 *a priori* echocardiographic hypotheses were conducted using a Bonferroni corrected *p*-value for significance (i.e., false discovery rate; FDR) of <0.0125.

Survival data were analyzed using the log-rank test; *p*<0.05 was considered significant. Between-groups differences were calculated using a 2-tailed student’s *t*-test for single paired comparisons where indicated; *p*-values <0.05 were considered significant. A Bayesian *p*-value was calculated for RNA-seq data as described previously.(3) Fold-change *p*-values of <0.001 were considered significant for RNA-seq data. For bioinformatics analyses, nominal p-values of <0.05 were considered significant for specific pathways or molecules of interest; *p*-values were also corrected for multiple hypothesis testing using the IPA software and a FDR <0.01 was considered significant. Please see the **Supplementary Methods** for more details on the statistical analyses provided by IPA software.

## RESULTS

### BET inhibition blunts negative cardiac remodeling and improves survival in PLN^R9C^

PLN^R9C^ mice display robust inflammatory and pro-fibrotic gene network activation before any phenotypic evidence of DCM (preDCM), but it is not clear if this plays a causal role in DCM progression.(3) BET inhibition has been recently shown to suppress this inflammatory program in other cardiomyopathy models.(13, 15) BET expression was robust in PLN^R9C^ hearts (**Supplementary Figure 3**). Therefore, we chose to target BETs in order to test if inhibition of this pathologic inflammatory gene expression program was a primary driver of DCM progression. We injected PLN^R9C^ or age-matched WT mice with the BET inhibitor JQ1 (**Figure 1A**). We first assessed the effects of BET inhibition on cardiac structure and function.

**Figure 1.**
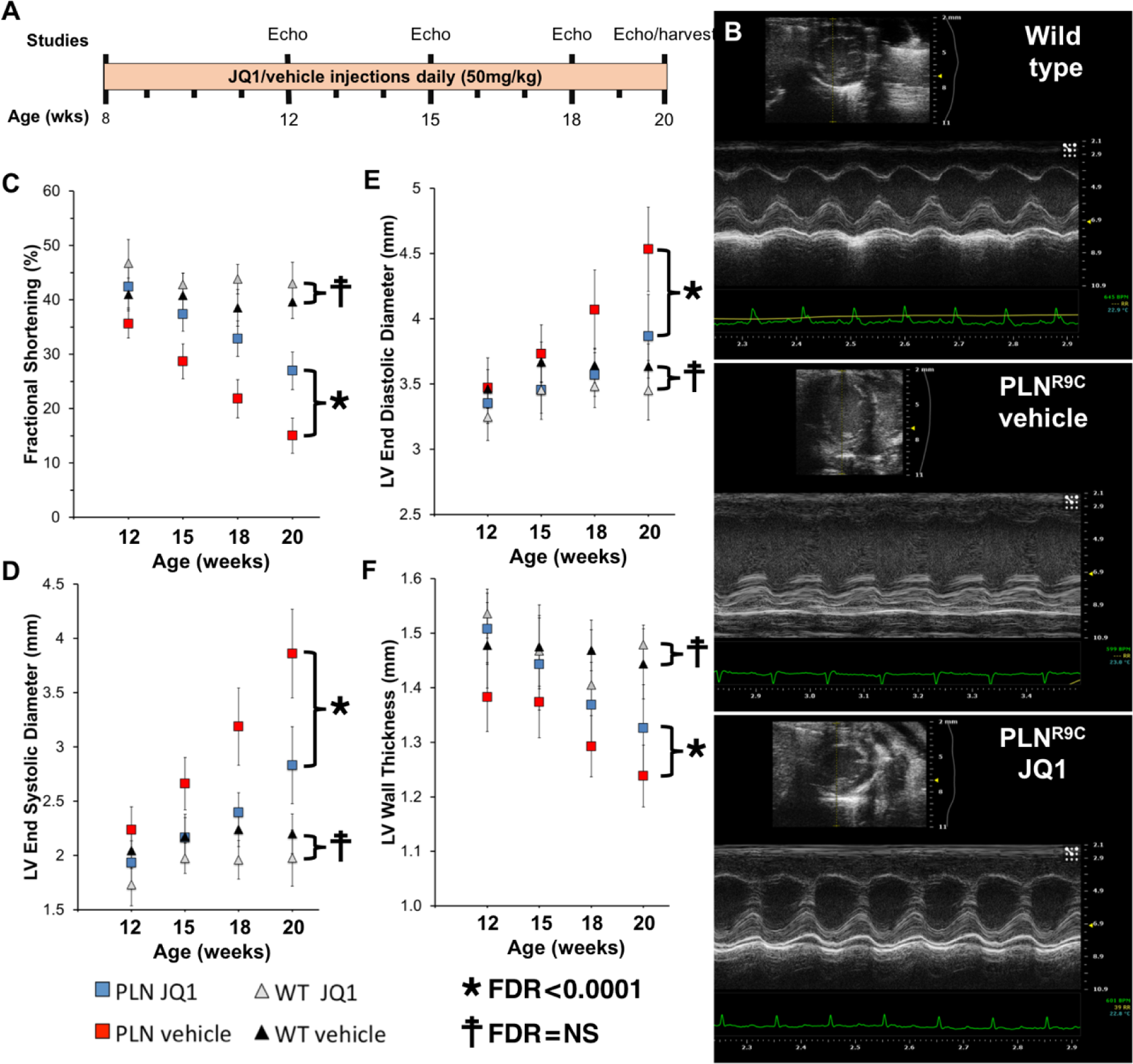
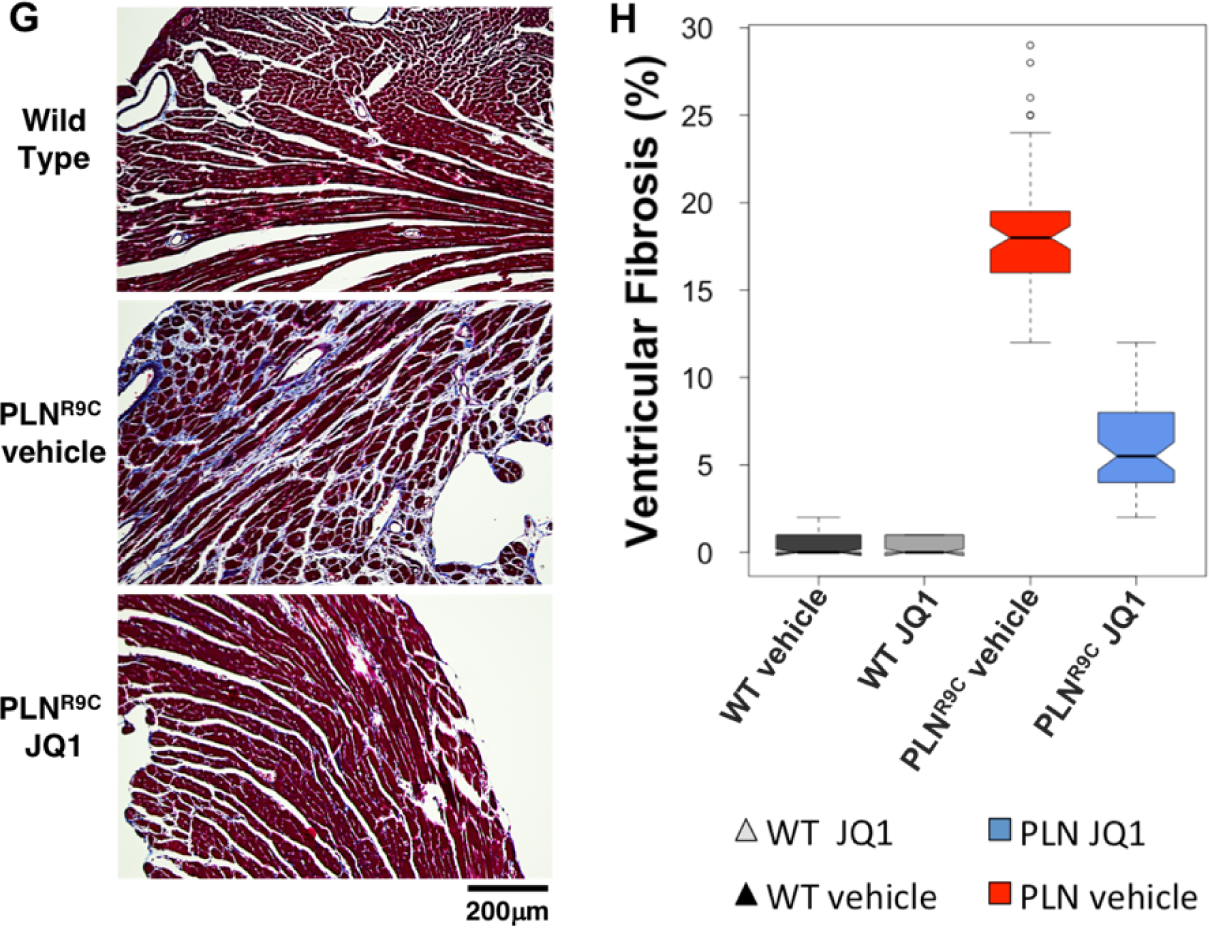
BET inhibition delays DCM in PLN^R9C^ mice. **(A)** Experimental protocol. **(B)** Representative M-mode images of WT and PLN^R9C^ vehicle or JQ1-treated mice at 18-weeks of age. **(C-F)** Echocardiographic assessment of mice treated with JQ1 or vehicle demonstrates progressive systolic dysfunction and negative left ventricular (LV) remodeling in PLN^R9C^ mice that was significantly blunted by JQ1 (n=14 PLN^R9C^, n=7 WT mice per group). **(G)** Representative Masson trichrome-stained LV sections from WT and PLN^R9C^ hearts. As JQ1 had no effect on fibrosis in WT, a single representative WT image is shown. **(H)** Quantification of scar area demonstrated severe fibrosis in PLN^R9C^ vehicle-treated hearts at 20-weeks of age that was markedly blunted by JQ1 (n=3 mice, 36 images from n=4 levels from apex to base for each mouse). Boxes = IQR; whiskers = 1.5x IQR; black line = median; notches = standard deviation, circles = extreme outlier values).

Treatment of WT mice with JQ1 had no effect on cardiac structure or function (**Figure 1**). PLN^R9C^ vehicle-treated mice demonstrated progressive negative cardiac remodeling as evidenced by reduced FS, increased LVESD, increased LVEDD, and decreased LVWT (p<0.001 for all) compared to WT vehicle-treated mice (**Figure 1**). By contrast, JQ1-treated PLN^R9C^ mice had better LV systolic function (smaller LVESD; higher FS; p<0.001), and less LV remodeling (smaller LVEDD; greater LVWT; p<0.001) than PLN^R9C^ vehicle-treated mice. Compared to WT vehicle-treated mice, PLN^R9C^ JQ1-treated mice still showed lower FS (p<0.001), but did not differ significantly from WT in LVESD, LVEDD or LVWT at 20-weeks of age. Similarly, heart weight to body weight ratio was significantly higher in PLN^R9C^-vehicle treated mice compared to WT (p<0.001), and returned to normal levels with JQ1 (**Supplementary Figure 4**).

PLN^R9C^ mice also develop tremendous fibrosis with progression to DCM.(3, 4) To assess if JQ1 could attenuate this response, hearts from 20-week old mice were stained with Masson trichrome and LV fibrosis quantified (**Figure 1G, H**). WT mice showed no fibrosis (0.3% of LV area). PLN^R9C^ vehicle-treated mice displayed severe fibrosis (18.4±0.7%; p<0.001 vs. WT vehicle-treated). JQ1 significantly reduced the amount of LV fibrosis by a factor of 3.2-fold (5.8±0.4%, p<0.001 vs. PLN^R9C^ vehicle-treated).

A separate cohort of mice (n=16) was followed for survival (**Figure 2**). Mice were treated with JQ1 or vehicle beginning at 8-weeks of age and monitored for mortality (as defined by our IACUC protocol). JQ1 treatment increased lifespan by 10% in PLN^R9C^ mice (p<0.001). These data demonstrate that JQ1 substantially delays disease progression and conveys a survival advantage in a robust model of chronic, progressive DCM.

**Figure 2.**
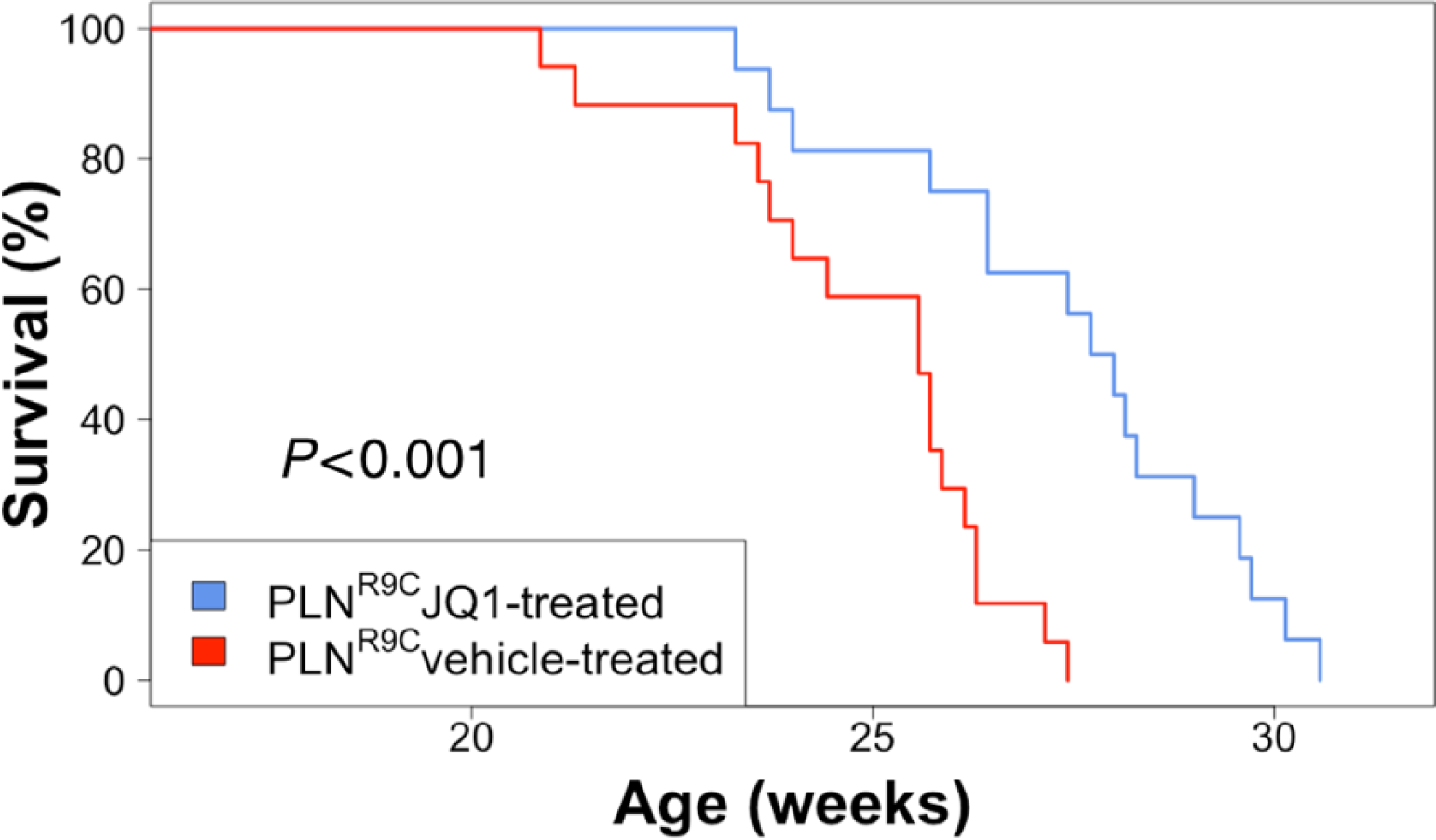
BET inhibition prolongs survival. Kaplan-Meier plots of survival free from death or severe HF symptoms.

### BETs control a pathologic gene expression program

Next, we assessed the primary transcriptional effects of BET inhibition at a very early stage of disease. We performed RNA-seq on whole LV tissue from PLN^R9C^ mice at the preDCM stage and age-matched WT controls treated with JQ1 or vehicle (**Figure 3A**, **Supplementary Table 1**). Principle components analysis (PCA) was used to confirm similar patterns of gene expression between mice in each treatment group. This demonstrated a pattern of gene expression in PLN^R9C^ vehicle-treated preDCM mice that was clearly distinct from the other groups, whereas JQ1 treatment normalized gene expression in PLN^R9C^ mice (**Supplementary Figure 5**). In WT mice, JQ1 triggered the differential expression of 692 genes (244 upregulated, 448 downregulated) across a wide spectrum of gene programs. After correction for multiple hypothesis testing, pathway analysis did not reveal enrichment of a concerted gene network suggesting that BETs regulate a subset of genes within many distinct gene programs at baseline (**Supplementary Table 2**).

**Figure 3.**
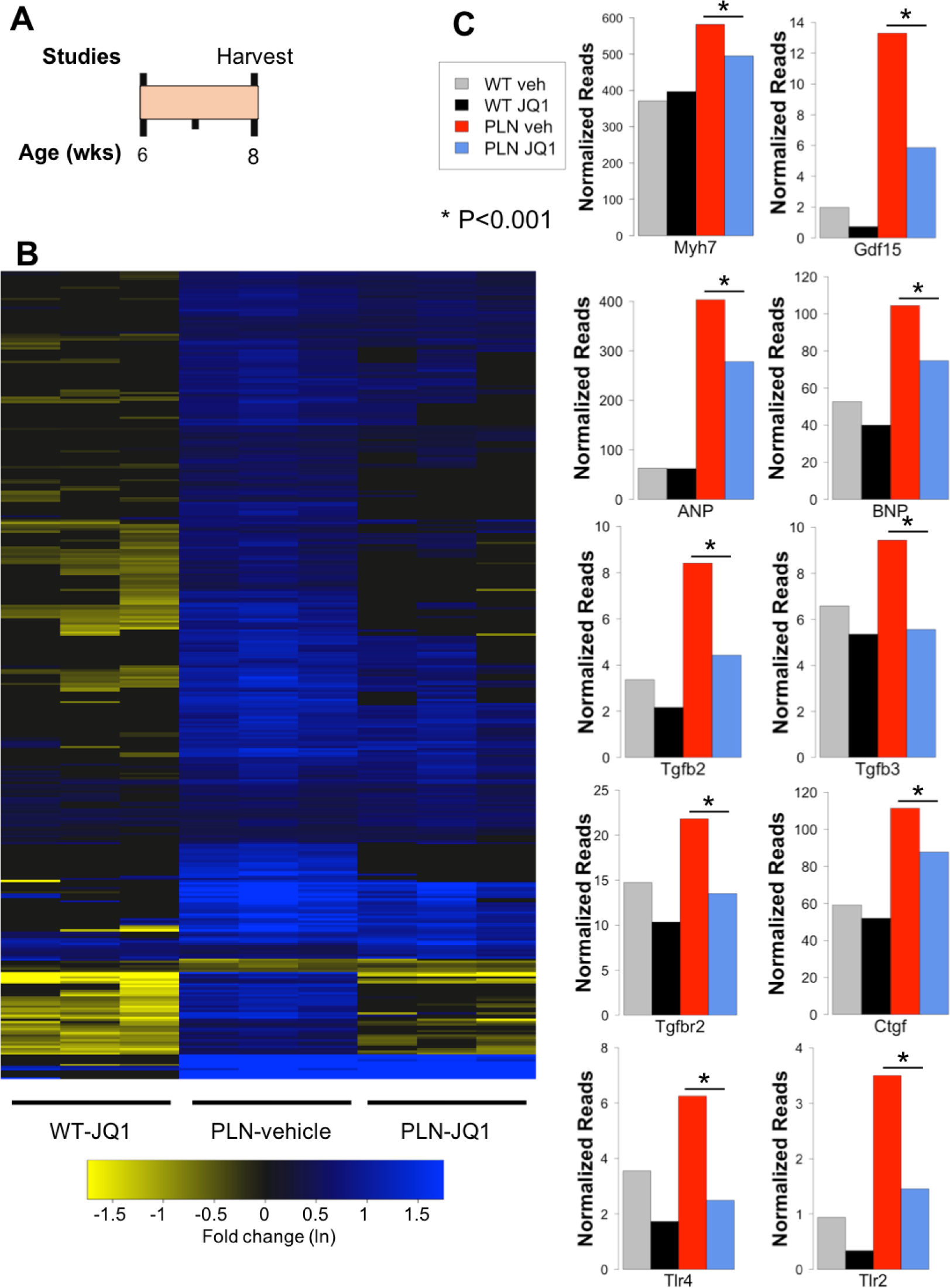
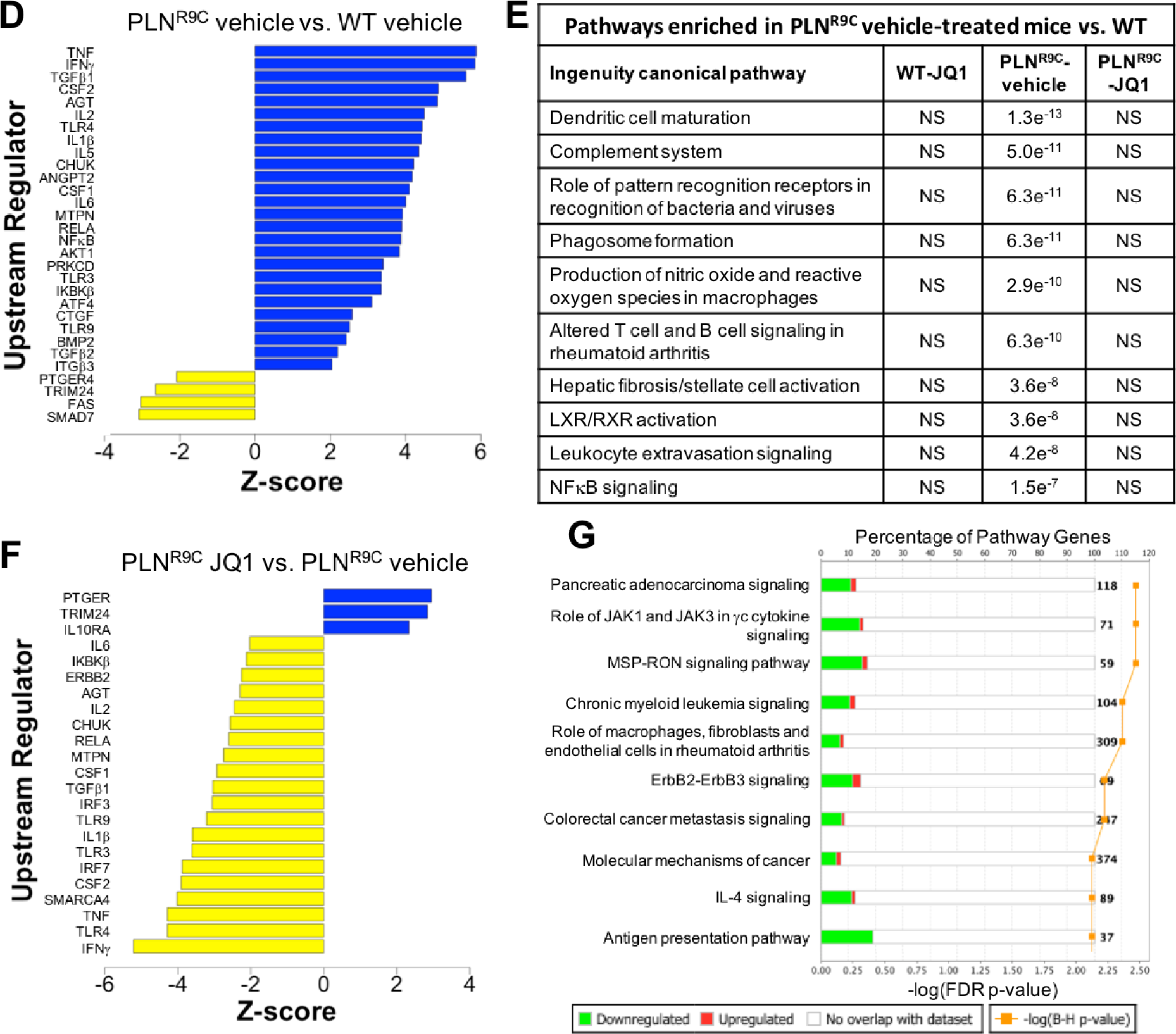
BETs activate inflammatory gene expression in preDCM hearts. **(A)** Experimental protocol. **(B)** Heat map of genes significantly differentially expressed in preDCM PLN^R9C^ vehicle-treated hearts demonstrating global reduction in gene activation in PLN^R9C^ JQ1-treated mice. Data plotted are natural log of fold-change values versus WT vehicle-treated mice. **(C)** Quantitation of key stress-response genes demonstrating marked upregulation in preDCM PLN^R9C^ hearts and significant reduction/normalization of mRNA levels with JQ1 treatment (n=3). **(D)** IPA upstream regulator analysis in preDCM PLN^R9C^ vehicle-treated mice compared to WT predicted the activation of many inflammatory mediators. **(E)** Selected IPA canonical pathways enriched in preDCM PLN^R9C^ vehicle-treated mice are no longer enriched in JQ1 treated hearts. Data listed are the false discovery rate (FDR) p-values. **(G)** IPA upstream regulator analysis demonstrating reduced activity of the majority of pro-inflammatory mediators in preDCM PLN^R9C^ JQ1-treated vs. PLN^R9C^ vehicle-treated mice. **(F)** Ingenuity canonical pathways from PLN^R9C^ JQ1-treated versus PLN^R9C^ vehicle-treated mice demonstrating suppressed gene expression within enriched pro-inflammatory pathways with JQ1 treatment (adapted from Ingenuity Pathway Analysis).

PreDCM PLN^R9C^ vehicle-treated hearts were heavily enriched for pro-inflammatory genes and pathways (**Supplementary Tables 1, 2**). Nearly all differentially expressed genes in PLN^R9C^ vehicle-treated hearts were upregulated (**Figure 3B**). Stress-response genes (growth-differentiation factor 15, the natriuretic peptides, fetal myosin heavy chain) were markedly increased in preDCM PLN^R9C^ vehicle-treated hearts as were genes involved in TGFβ signaling (*Tgfb2, Tgfb3, Tgfbr2, Ctgf*) and the innate immune response (*Tlr2, Tlr4*; **Figure 3C**). The pattern of differentially expressed genes in PLN^R9C^ vehicle-treated mice predicted activation of key inflammatory signaling molecules and transcriptional regulators, implicating TGFβ, NFκB, AKT and both the innate and adaptive immune response systems as major early effectors of DCM (**Figure 3D, E**, **Supplementary Table 3**).

Treatment with JQ1 strongly attenuated these gene expression changes (**Figure 3B, C**). Relative to vehicle-treated PLN^R9C^ mice, JQ1 reduced the expression of 663 genes (80.1% of differentially expressed genes). Pathway analysis revealed near uniform suppression of inflammatory gene networks and pro-inflammatory upstream regulators (i.e., TGFβ, TLRs, and a variety of cytokines) by JQ1 in preDCM PLN^R9C^ hearts (**Figure 3E-G**). These data demonstrate that BETs strongly and specifically regulate inflammatory gene network activation in preDCM LV tissue, a stage where changes in LV geometry, systolic function or fibrosis have yet to manifest.

### Chronic BET inhibition dampens cardiac inflammatory signaling

To study the effects of long-term BET inhibition, mice were treated from 8- to 20-weeks of age with JQ1 or vehicle and RNA-seq performed (**Figure 4A**). PCA again demonstrated excellent grouping of mice, and a shift towards WT gene expression in PLN^R9C^ JQ1-treated hearts (**Supplementary Figure 6**). PLN^R9C^ vehicle-treated mice with overt DCM demonstrated progressive inflammatory gene network activation (**Supplementary Tables 4, 5**) concomitant with a shift in myocardial metabolic gene expression; we found near uniform decreases in aerobic metabolic genes including those involved in mitochondrial oxidative phosphorylation, fatty acid oxidation and tricarboxylic acid (TCA) cycle (**Figure 4B**), with concurrent increases in expression of genes controlling cellular glucose utilization (**Figure 4C**).

**Figure 4:**
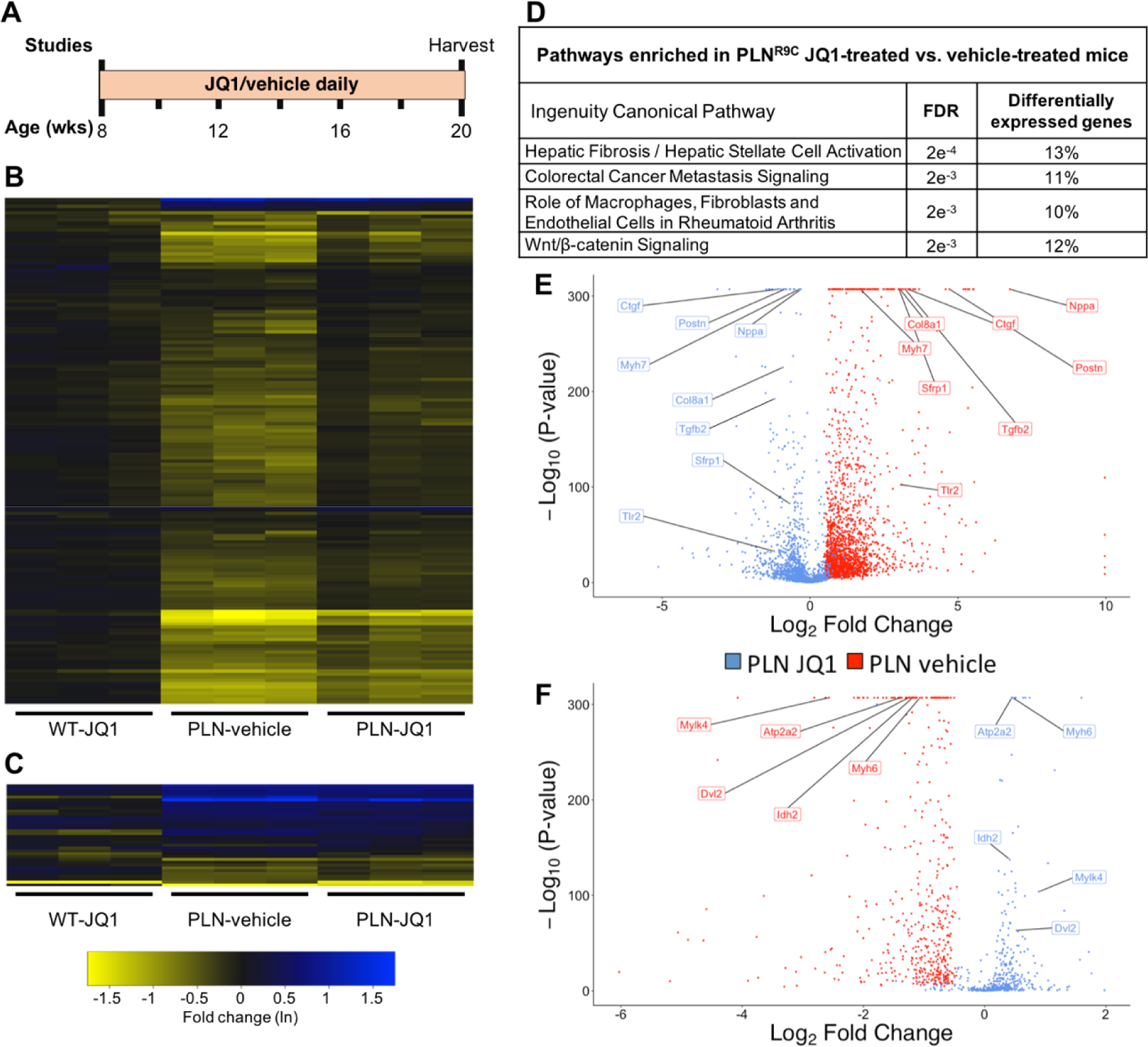
Chronic BET inhibition alters inflammatory gene expression in DCM. **(A)** Experimental protocol. PLN^R9C^ mice with overt DCM displayed a profound shift in metabolic gene expression. Heat maps demonstrating **(B)** genes controlling aerobic respiration (tricarboxylic acid cycle and mitochondrial oxidative phosphorylation) were almost uniformly downregulated, while **(C)** genes controlling cellular glucose utilization were generally upregulated in PLN^R9C^ hearts. JQ1 had little effect on metabolic gene expression. **(D)** Enriched pathways between PLN^R9C^ JQ1-treated and vehicle-treated mice. Differentially expressed genes are the percent of pathway genes differentially expressed between JQ1 and vehicle treated PLN^R9C^ mice. Volcano plots demonstrating the magnitude and significance of JQ1 on altering the expression of genes that were **(E)** upregulated or **(F)** downregulated in PLN^R9C^ vehicle-treated hearts. In each plot, blue represents the effect of JQ1 on gene expression for all genes in red. Representative genes (labeled) reveal key processes affected by JQ1 including cardiac stress response signaling (*Nppa*, *Myh6*/*Myh7*), fibrosis and extracellular matrix remodeling (*Postn*, *Col8a1*), TGFβ signaling (*Tgfb2*, *Ctgf*), cytoskeletal signaling (*Mylk4*), Wnt signaling (*Sfrp1*, *Dvl2*), innate immune activation (*Tlr2*, *Tlr4*), and metabolism (*Idh2*, *Atp2a2*).

Longitudinal treatment with JQ1 continued to suppress pathologic inflammatory gene expression, though the effect on metabolic gene expression was only partial at this late stage of disease. Pathways suppressed (**Figure 4D**) included fibrosis and Wnt-signaling networks. While many genes and pathways are still dysregulated in PLN^R9C^ JQ1-treated mice, the magnitude of effect remained substantially lower than that observed in PLN^R9C^ vehicle-treated mice, demonstrating the continued effect of JQ1 in late stage DCM (**Figure 4E, F**). Collectively, these data show that BET inhibition in preDCM mice potently and specifically suppresses pathologic inflammatory and pro-fibrotic gene networks, substantially reduced myocardial fibrosis and blunts negative cardiac remodeling, thus establishing a causal role for inflammatory gene networks in DCM progression.

### BET inhibition suppresses fibroblast activation

Given the salutary effects on inflammatory gene network expression and fibrosis, we tested whether BET inhibition had differential effects in the different cardiac cell compartments. We previously showed excess non-myocyte proliferation in PLN^R9C^ mice.(3) To test if BETs play a role in the proliferation of non-myocytes in this model, PLN^R9C^ and WT mice treated with JQ1 or vehicle were treated with EdU prior to sacrifice (**Figure 5A**). Proliferating non-myocytes were 3.7-fold more prevalent (p<0.001) in PLN^R9C^ vehicle-treated mice compared to WT (**Figure 5B, C**). This was reduced 1.9-fold by JQ1 (p<0.001). Additionally, our RNA-seq data showed a strong increase in genes that are characteristic of myofibroblasts in DCM hearts; these were reduced by JQ1 (**Supplementary Figure 7**). Therefore, we hypothesized that these proliferating cells were cardiac fibroblasts.

**Figure 5:**
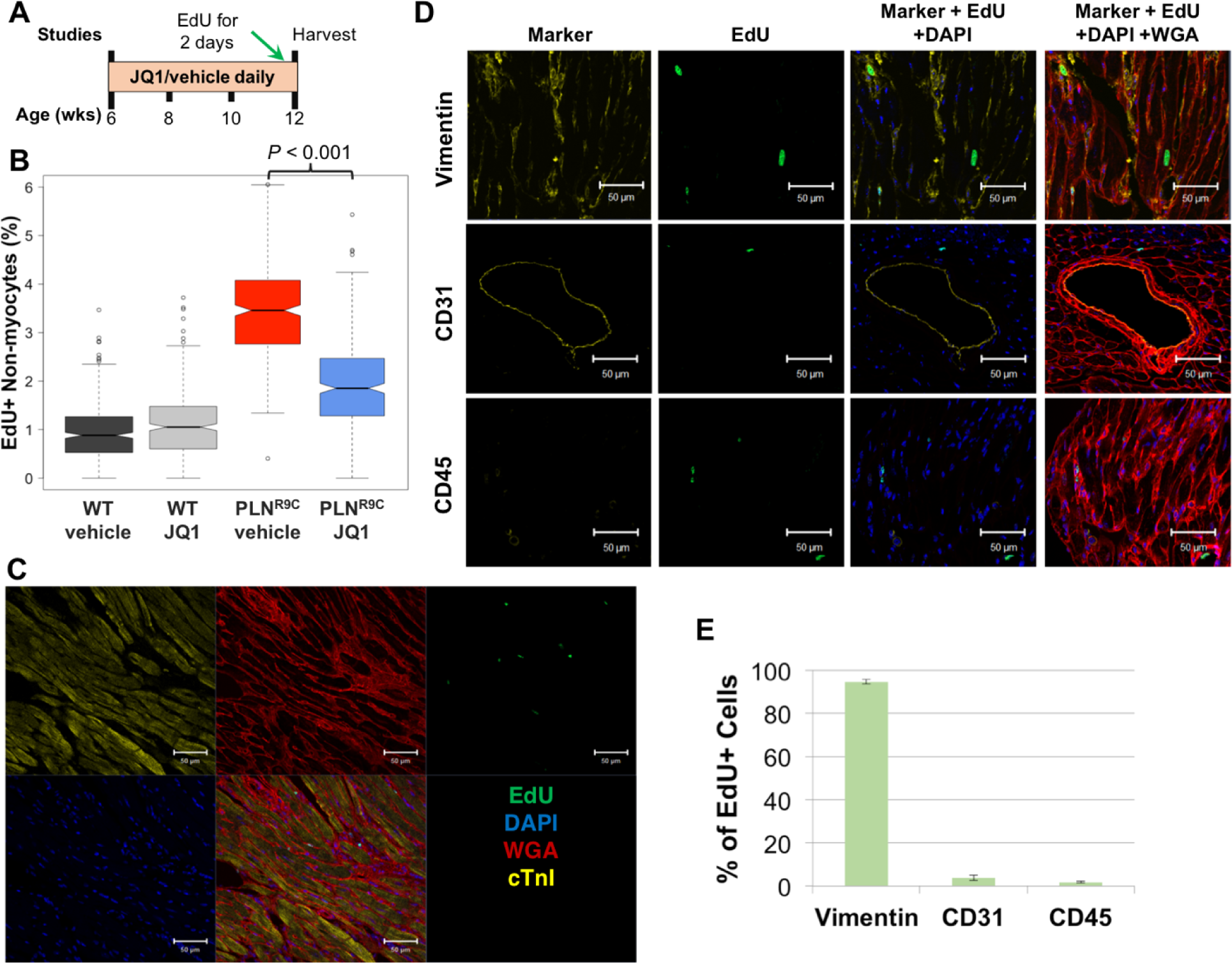
BET inhibition prevents cardiac fibroblast proliferation. **(A)** Experimental protocol. **(B)** PLN^R9C^ vehicle-treated mice displayed 3.7-fold more proliferating non-myocytes than WT mice. This was substantially reduced by JQ1. Boxes display the interquartile range (IQR), notches show the SD, whiskers are 1.5x the IQR and circles are extreme outliers. **(C)** Representative confocal image showing images stained for EdU, DAPI, WGA, cardiac Troponin I (cTnI), and merged. **(D)** Representative images from a multi-marker analysis demonstrating co-staining of EdU with vimentin but not with CD31 or CD45. **(E)** Quantification of EdU-positive, marker-positive cells identified the majority of EdU cells as cardiac fibroblasts. Slides were co-stained with DAPI and WGA.

As there is no single consensus fibroblast marker, we undertook a multi-marker histochemical approach to identify the nature of the EdU-positive cells in PLN^R9C^ hearts. Vimentin staining in the absence of co-staining for CD31 (endothelial cells) and CD45 (myeloid cells) is an accepted means of identifying fibroblasts.(18, 19) We found the vast majority (94.6±1.0%) of EdU-positive non-myocytes co-stained with vimentin, with only a small percentage staining with CD31 (3.7±1.2%) or CD45 (1.8±0.4%), thus confirming that the proliferating cells were predominantly cardiac fibroblasts (**Figure 5D, E**). There was no significant difference in the percentage of EdU-positive cells co-stained with vimentin, CD31 or CD45 in WT or PLN^R9C^ mice treated with either vehicle or JQ1 (data not shown). We also observed significant decreases in genes associated with myofibroblast activation in DCM mice treated with JQ1, including *Postn* (-1.9-fold, p<0.001), *Eln* (-3.0-fold, p<0.001), *Lox* (-1.7-fold, p<0.001), *Mfap2* (-1.4-fold, p<0.001), *Itga1* (-1.5-fold, p<0.001) and *Pdgfra* (-1.6-fold, p<0.001). These data affirm the role of BETs in cardiac fibroblast activation in DCM.

Next, to test the cell compartment specific effects of JQ1 on gene expression, we performed RNA-seq on isolated pools of cardiac non-myocytes and cardiomyocytes (**Figure 6**). JQ1 had a substantial effect on the pattern of gene expression in non-myocytes (**Figure 6B**), but not cardiomyocytes (**Figure 6E**). Similarly, JQ1 drove a significant change in gene expression in PLN^R9C^ non-myocytes (**Figure 6D**) but not cardiomyocytes (**Figure 6F**). Enriched pathways in PLN^R9C^ non-myocytes predominantly consisted of inflammatory gene networks that were activated with progression to DCM, an effect completely eliminated by JQ1 (**Figure 6C**). By contrast, cellular metabolic pathways (the predominant program altered in DCM cardiomyocytes) all remained enriched in PLN^R9C^ cardiomyocytes after JQ1 treatment (**Figure 6G**). Collectively, these data demonstrate that BETs play a critical role in the expression of dynamically regulated inflammatory gene networks in cardiac non-myocytes, and suggest that BET inhibition does not exert its salutary effects on heart function in this model via modulation of adult cardiomyocyte gene expression programs.

**Figure 6:**
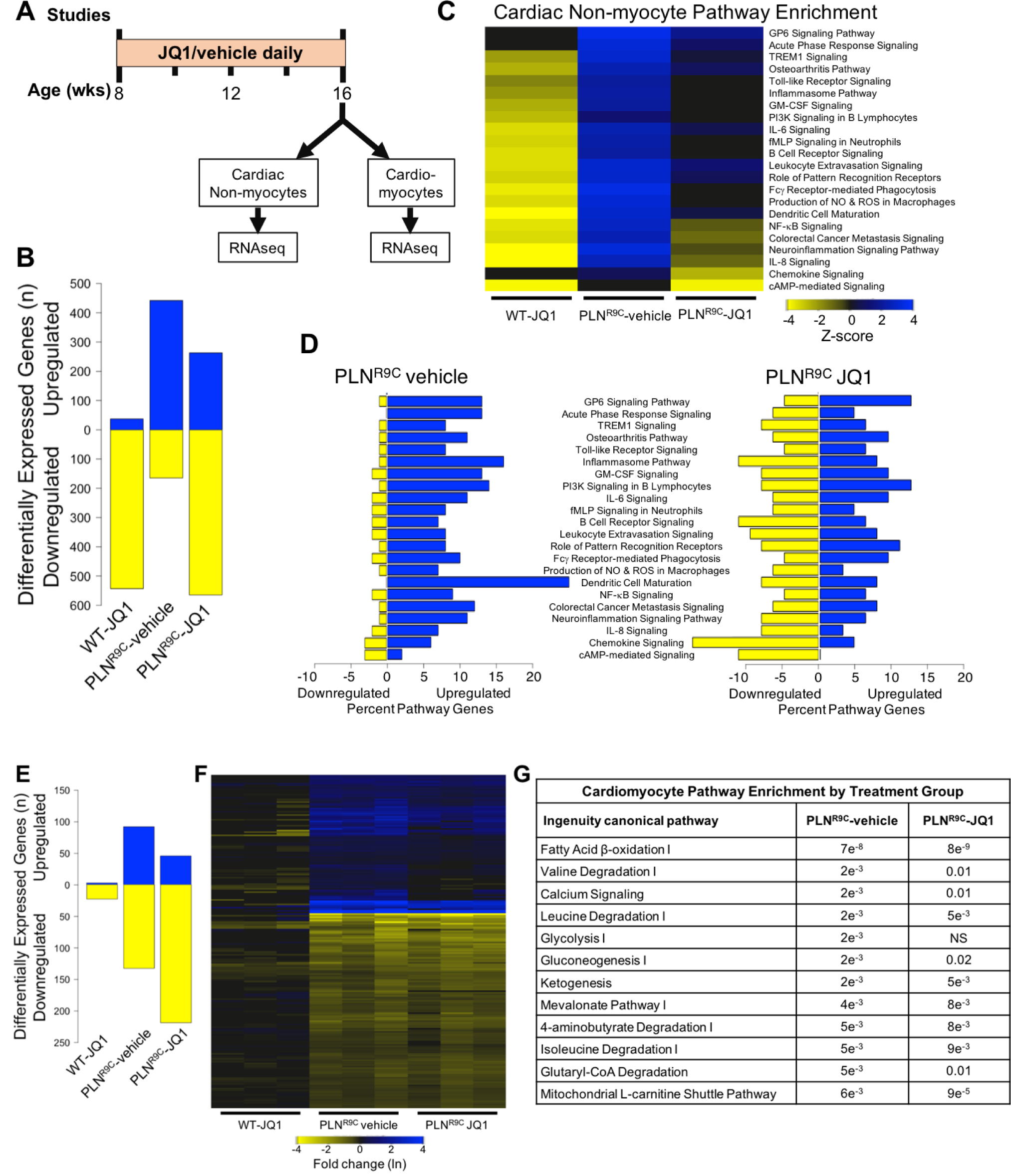
BETs primarily affect cardiac non-myocyte gene expression programs. **(A)** Experimental protocol. **(B)** Differentially expressed genes from pooled cardiac non-myocytes (relative to WT vehicle-treated mice) were predominantly downregulated in WT JQ1-treated mice and upregulated in PLN^R9C^ vehicle-treated mice. Non-myocytes from PLN^R9C^ mice treated with JQ1 displayed a marked shift in gene expression. **(C)** Pathways enriched in non-myocytes from PLN^R9C^ vehicle-treated mice with z-score ±2 were all downregulated or not enriched in mice treated with JQ1. **(D)** JQ1 treatment shifted gene expression in those pathways from predominantly up to predominantly downregulated. **(E)** The pattern of differentially expressed genes from pooled cardiomyocytes (relative to WT vehicle-treated mice) was similar between PLN^R9C^ vehicle-treated and JQ1-treated mice. Further, JQ1 had virtually no effect on gene expression in WT cardiomyocytes. **(F)** Heat map of genes differentially expressed in PLN^R9C^ vehicle-treated cardiomyocytes showing little to no change in expression levels with JQ1 treatment. **(G)** Pathway enrichment in PLN^R9C^ cardiomyocytes was dominated by metabolic pathways, which were uniformly still enriched in PLN^R9C^ JQ1-treated cardiomyocytes.

### BETs interact with NFκB in cardiac fibroblasts

The mechanisms by which BETs preferentially suppress inflammatory gene expression in this model of chronic DCM remain unknown. BRD4, the archetypal BET, is necessary for NFκB-mediated inflammatory gene expression in models of atherosclerosis,(20) graft-versus-host disease,(21) and cancer.(22–24) Thus, we investigated if BETs exert their potent effects on non-myocyte gene expression via NFκB. Our extensive RNA-seq studies revealed that NFκB signaling was strongly upregulated in PLN^R9C^ hearts and isolated cardiac non-myocytes at all disease stages, and genes activated by NFκB were strongly upregulated in cardiac non-myocytes (**Figure 7A, B**).

**Figure 7:**
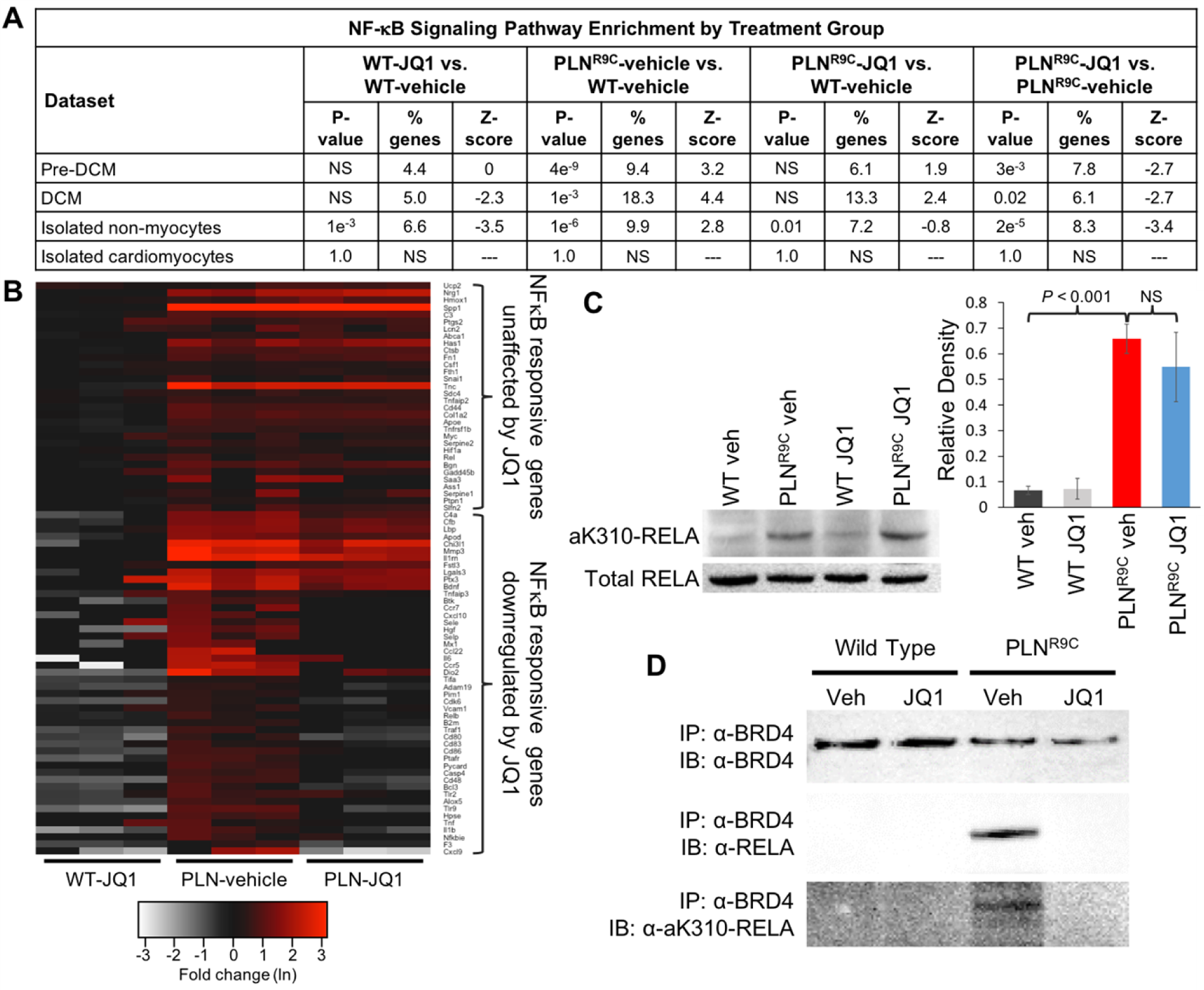
NFκB is activated in a BRD4-dependent manner in PLN^R9C^ hearts. **(A)** The Ingenuity pathway NFκB signaling was highly enriched in hearts from preDCM and DCM PLN^R9C^ mice, but not WT, and was specifically enriched in the cardiac non-myocyte cell population. This association was prevented by BET inhibition, with strong downregulation of the pathway by JQ1 therapy. **(B)** Heat map of genes known to be activated by NFκB demonstrated uniform upregulation in cardiac non-myocytes with a majority of genes being switched off by JQ1 therapy. **(C)** Western blot showing increased RELA acetyl lysine-310 (aK310) in PLN^R9C^ hearts as compared to WT irrespective of JQ1 treatment status (representative images of n=3 experiments). **(D)** Western blots (IB) from the co-immunoprecipitation (IP) of BRD4 with anti-BRD4, anti-RELA and anti-ak310-RELA demonstrated direct binding of BRD4 and NFκB in PLN^R9C^ hearts with DCM. This was completely abolished by treatment with JQ1 (representative images of n=3 experiments).

Acetylation of the RELA subunit of NFκB is a well-established marker of increased NFκB activity.(25) We identified markedly increased aK310-RELA in PLN^R9C^ hearts with DCM (**Supplementary Figure 8**). This finding was independent of JQ1 treatment (**Figure 7C**), suggesting NFκB activation occurs upstream of BET action as was previously demonstrated in endothelial cells.(20)

Next, we performed co-immunoprecipitation (IP) of BRD4 with RELA and aK310-RELA, demonstrating a clear physical association in PLN^R9C^ mice with DCM, an association that was abolished by treatment with JQ1 (**Figure 7D**). IP of RELA confirmed the binding of BRD4 with NFκB (data not shown). These data reveal that in this model of DCM, BET-mediated gene expression occurs, at least in part, via co-activation of NFκB.

## DISCUSSION

We have demonstrated that BET bromodomain inhibition is an effective strategy for disrupting the progression of chronic non-ischemic HF. Our comprehensive transcriptomic analyses revealed that BETs control a pathologic gene expression program that is dominated by inflammatory and profibrotic signaling networks in the heart. Longitudinal treatment with JQ1 delayed the onset and progression of DCM, confirming that this inflammatory milieu plays a central role in DCM progression. Importantly, we discovered that BET inhibition has profound and specific effects on fibroblast activation and gene expression with minimal concomitant effects in cardiomyocytes. Finally, we identified an important mechanistic link between the BET family member BRD4 and NFκB in cardiac fibroblasts. Collectively, these data highlight the central role of BETs in DCM pathobiology and define a critical role for BETs in cardiac fibroblast activation. These data confirm BET inhibition and transcriptional regulation as a potential novel therapeutic strategy for HF.

BET inhibition with JQ1 has been shown to reduce myocardial hypertrophy and fibrosis induced by phenylephrine or thoracic aortic banding, and limits infarct size and HF after myocardial infarction. (13–15) These studies have collectively demonstrated potent suppression of gene networks involved in inflammation, fibrosis and hypertrophic growth. We have extended these findings to a model of genetic DCM. The PLN^R9C^ mouse develops slowly progressive disease in early-adult life. Importantly, in contrast to preclinical models of acute cardiac stress, PLN^R9C^ is a model of insidiously progressive DCM that mimics the tempo of HF that is typically seen in humans. BET inhibition with JQ1 slowed the progression of DCM, prevented fibrosis and conferred a survival advantage in PLN^R9C^ mice. These findings are particularly relevant to human HF, a disease process that, despite intensive research for decades, still leaves the vast majority of individuals with some degree of functional limitation and carries a 5-year mortality in excess of 50%.(1) Astoundingly, current HF pharmacotherapy (neurohumoral blockade, selected vasodilator regimens and now possibly SGLT-2 inhibitors) is solely based on our understanding of the systemic consequences of HF. New strategies, particularly those focused on disrupting intrinsic disease-specific myocardial processes are urgently needed.

We have previously shown early and progressive activation of inflammatory gene networks in the PLN^R9C^ non-myocyte population.(3) We have now demonstrated that BETs are central to inflammatory gene network expression in DCM and, importantly, that this effect occurs selectively in the cardiac fibroblast population. Recent evidence has shown a central role for BRD4 in rat cardiac fibroblast activation.(26) Our data also revealed that BETs have a direct role in fibroblast proliferation and myofibroblast activation. As myofibroblasts in the heart are almost exclusively derived from resident cardiac fibroblasts,(19) we conclude that BETs play a central role in the transdifferentiation of resident cardiac fibroblasts to myofibroblasts in PLN^R9C^ hearts. Thus, we have confirmed BETs as nodal mediators of fibroblast activation in chronic DCM.

Interestingly, this evolving inflammatory milieu precedes fibrosis, LV remodeling and overt cardiomyocyte pathology in spite of the fact that PLN^R9C^ is a disease caused by a cardiomyocyte-specific gene mutation that alters cardiomyocyte calcium handling.(4) Thus, the inflammatory gene network activation in PLN^R9C^ hearts is a secondary consequence of early cardiomyocyte changes that is triggered by an as-yet undefined mechanism. This left open the question of whether these secondary inflammatory changes were truly a driver of DCM progression or an epiphenomenon. By targeting BET bromodomains with JQ1, we were able to specifically and selectively block inflammatory gene network activation, which substantially inhibited DCM progression, thereby demonstrating a causal role for inflammatory gene network activation in DCM progression. This finding also highlights the potential therapeutic benefit of a strategy targeting transcription in HF.

To extend these findings, we also performed RNA-seq on pooled cardiomyocytes. Interestingly, in WT mice, there were virtually no gene expression changes in response to JQ1. Further, while some gene expression changes were evident in cardiomyocytes from PLN^R9C^ JQ1-treated mice, the vast majority of genes differentially expressed were not substantially affected by JQ1. By way of example, the predominant gene expression programs affected in DCM cardiomyocytes are metabolic networks, an effect we previously attributed to downregulation of peroxisome-proliferator activated receptor (PPAR) signaling.(3) We confirmed this shift in metabolic gene expression from PLN^R9C^ vehicle-treated cardiomyocytes. Yet, we observed minimal changes in global cardiomyocyte metabolic gene expression and specifically saw minimal differences in PPAR network genes, which largely remained downregulated, in spite of JQ1 therapy. Thus, we conclude that BETs limit cardiac remodeling primarily by blocking the pathologic inflammatory gene program in cardiac fibroblasts with little concomitant effect on gene transcription in cardiomyocytes.

Previous work using cultured neonatal rat ventricular cardiomyocytes treated with phenylephrine, and human induced pluripotent stem cell-derived cardiomyocytes (a cell type that assumes an immature gene expression profile upon differentiation) treated with endothelin-1 demonstrated dynamic changes in cardiomyocyte gene expression.(15, 26) However, in vivo gene signatures from these same studies were still dominated by inflammatory and myofibroblast-specific gene programs. Our ex vivo analysis is, to our knowledge, the first to isolate cardiomyocyte and non-myocyte cell fractions and shows that the dominant gene expression signature is in the cardiac fibroblasts. Interestingly, physiologic hypertrophy triggered by high-intensity swimming, which is not associated with inflammatory network activation or a fibrotic response, was not affected by JQ1.(15) Perhaps this is due to the limited role of BETs on normal gene expression in adult cardiomyocytes. Similarly, it may be possible that BETs have a more prominent role in embryonic and/or neonatal cardiomyocytes. Further studies are needed to clarify the role of BETs in adult cardiomyocytes.

The mechanism by which BETs exert their effect on inflammatory gene transcription in cardiac fibroblasts is steadily taking shape, though the picture remains incomplete. BETs bind to acetylated lysine residues on histone tails at promoters of active genes. The binding of BRD4, specifically, recruits the positive transcription elongation factor-b complex, which promotes pause release of RNA polymerase II, facilitating gene transcription.(13, 27, 28) Numerous studies have shown that BRD4 is found at enhancer sequences genome-wide, including in the heart.(13, 29) However, under stress conditions, BRD4 is rapidly redistributed to super enhancers (SEs), driving disease-specific gene expression. SEs are cis-regulatory elements that permit dense clustering of DNA-bound transcription factors that are associated with exceedingly high levels of transcriptional co-activators, including BRD4.(30, 31) SE activation alters the cellular transcriptome and can permit cell-state transition.(20, 32) In the heart, this manifests as pathologic gene expression and the transdifferentiation of quiescent fibroblasts to myofibroblasts.(29)

Precisely how this locus-specific flux of BRD4 to DCM-specific SEs occurs has not previously been elucidated in the heart. For the first time in cardiac fibroblasts, we have shown a direct mechanistic link between BRD4 and acetylated RELA, demonstrating that BRD4 exerts its effects on gene expression, at least in part, via NFκB. BRD4 has been shown to bind acetylated lysine residues on the p65/RelA subunit of NFκB, thus driving gene expression.(22, 23, 33) In endothelial cells, cytokine treatment triggers reorganization of chromatin activators from basal endothelial cell enhancers to inflammatory SEs, triggering inflammatory gene expression via NFκB in a BRD4-dependent manner.(34) Importantly, these authors very elegantly demonstrated that NFκB activation and SE binding appears to occur upstream of BRD4 as NFκB was still bound to SEs in the absence of BRD4. Hence, JQ1 disrupts the ability of NFκB to recruit BRD4 to stress-activated SEs, thus limiting changes in transcription. This likely explains why we did not see a significant decrease in aK310-RELA in PLN^R9C^ JQ1-treated hearts relative to vehicle. Future research should be aimed at characterizing the upstream activators of NFκB and identifying additional cardiac transcription factors that may recruit BRD4 to SEs in activated cardiac fibroblasts.

Our data adds an important element to the BET story in the heart, yet many important questions remain. While this study contributes to mounting evidence that highlights the clear importance of BRD4 in pathologic cardiac gene expression, the roles of BRD2 and BRD3 in the heart have not been defined. In cultured rat cardiac fibroblasts, *Brd2* inhibition did result in reduced expression of myofibroblast markers in response to cytokine stimulation, suggesting BRD2 has some role in fibroblast gene expression.(29) Further, in immune cells, BRD2 binds to the chromatin insulator CTCF; this facilitates SE formation, which in turn recruits BRD4, driving gene expression and cell-state transitions via coordinate BRD2-BRD4 interactions at these newly commissioned SEs.(35) Whether such a role exists for BRD2 in cardiac fibroblasts remains unknown. Little is known of the biologic function of BRD3. Additional questions also surround the role of cardiomyocyte expressed BET isoforms.

This study has several limitations. First, mRNA transcript levels do not always correlate with protein levels or protein activity and are thus an indirect measure of pathway activation in vivo. Despite normalization of RNA libraries, the demonstrated proliferation of cardiac fibroblasts confounds the precise interpretation of gene transcript changes among non-myocyte genes in bulk RNA-seq experiments from whole LV samples. Finally, pathway analysis tools are inherently limited by the existing knowledge base for a given gene.

Fundamentally, HF is a disease of dysregulated gene expression, much like cancer. However, the mechanisms are distinct: in cancer, the progressive accumulation of mutations drives pathologic gene transcription that ultimately lead to dysregulated cell growth; whereas in HF, the continuous activation of stress signaling networks converges on the transcriptional machinery to alter the myocardial cellular phenotype, ultimately hampering cardiac function. Current HF treatments that target these systemic stress-signaling networks are effective, but morbidity remains exceedingly high. Consequently, there is an urgent unmet clinical need to directly target intramyocardial processes to improve outcomes.(36) Upstream stress signaling networks converge on BET bromodomain proteins, which integrate these signals into specific and distinct gene expression programs. As such, BET inhibition is a particularly appealing therapeutic strategy that could be both additive to neurohumoral blockade and widely applicable to HF of varying etiologies. These data have significantly extended our understanding of cardiac BET biology and proved the effectiveness of BET inhibition in a model of chronic DCM that closely mimics human HF. Concurrent with ongoing trials in cancer, this knowledge should serve as a springboard for advancing BET and other chromatin-based therapeutic strategies for the treatment of HF.

