## Supplementary Materials for "BET Bromodomain Proteins Regulate Transcriptional Reprogramming in Genetic Dilated Cardiomyopathy"

### SUPPLEMENTARY METHODS

*Tissue Harvesting:* $PLN^{R9C}$ or age-matched WT mice were deeply anesthetized with isoflurane; hearts were exposed by midline thoracotomy and immediately excised. Hearts were rinsed in cold 1x PBS to remove blood and then subjected to ventriculectomy (atria, atrioventricular valves and residual vascular tissue were removed) followed immediately by either tissue fixation or flash freezing for subsequent RNA or protein extraction. For frozen tissue, the right ventricle was also excised.

*Histochemistry and quantification of myocardial fibrosis:* Hearts were fixed in 4% paraformaldehyde, paraffin-embedded and sections cut from apex to base to cover all regions of the ventricular tissue. Sections at multiple levels from apex to base (up to 15 as ventricular size permitted) were stained with Masson trichrome. Each section contained 6 pieces of ventricular tissue in book-matched pairs, permitting analysis of up to 90 distinct pieces of LV tissue per mouse.

Fibrosis was quantified as the ratio of fibrotic area to total ventricular area using a Keyence BZ-X700 microscope. Briefly, a section of ventricular tissue was scanned at 10x magnification. Tiled images were stitched together (automatically performed by the Keyence BZ-X700 software) and blue hues, corresponding to collagen deposition in Masson trichrome stained slides, were identified and quantified by the microscope software as a ratio relative to the remaining ventricular tissue.

*Immunohistochemistry:* Hearts were fixed and paraffin-embedded as described above. For all immunohistochemical experiments, sections were deparaffinized with xylenes and ethanol, rehydrated in deionized water and PBS, and then permeabilized with 0.1% Tween in PBS. Antigen retrieval was performed by heating slides in target retrieval solution (Diva Decloaker, Biocare Medical) at 125°C for 30 minutes.

To identify proliferating cells, we utilized 5-ethynyl-2'-deoxyuracil (EdU, Thermo-Fisher).(1) Mice were injected IP with EdU on 2 consecutive days prior to sacrifice. Twenty four hours after the final EdU injection, hearts were harvested. To detect EdU, slides were stained using the Click-iT EdU Alexa Fluor 488 Imaging Kit (Thermo-Fisher). Sections were blocked (10% normal goat serum, 1% BSA, 0.1% Triton X-100 in PBS) for 30 minutes at room temperature. Next, slides were stained with 1:200 anti-wheat germ agglutinin (WGA) Alexa Fluor 555 (Molecular Probes, #W32464) in 0.05% Tween-20, 1X PBS for 30 minutes at 37°C, then washed in 1X PBS-T. Finally, slides were stained with DAPI (Sigma-Aldrich, #D9542), washed with 1X PBS and then mounted with VectaShield (Vector labs).

Microscopy was performed using a Zeiss LSM510 confocal microscope. To maximize the number of cells counted, EdU-labeled cells were counted manually at 10x magnification from 24 fields/section at 7 different levels from apex to base per animal (n=3 mice/group); >70,000 nuclei were counted per animal. Fields were chosen at random from different regions of the ventricle to minimize selection bias prior to fluorescence imaging. DAPI-labeled cells were counted at 10x magnification using Image J.

To our knowledge, the ideal amount of EdU for cardiac studies is not known, though studies in brain tissue have suggested saturation of proliferating cells at a dose of 50mg/kg.(2) Thus, we first injected mice with EdU at doses of 10mg/kg, 25mg/kg and 50mg/kg (n=3 WT mice each dose). As shown in **Supplementary Figure 1**, we chose to proceed with EdU at a dose of 25mg/kg. Hence, after 6-weeks of treatment with JQ1 or vehicle, $PLN^{R9C}$ mice and age-matched controls were injected IP on 2 consecutive days prior to sacrifice with EdU at 25g/kg body weight in 10% DMSO-PBS at 14-weeks of age. Mice were sacrificed approximately 24 hours after the last injection. Hearts were harvested, fixed, and processed as above. After rehydration, slides were stained using the Click-iT EdU Alexa Fluor 488 Imaging Kit. Sections were then blocked (10% normal goat serum, 1% BSA, 0.1% Triton X-100 in PBS) for 30 minutes at room temperature, followed by staining with 1:250 anti-cardiac troponin I (cTnI) antibody in 3% BSA-1X PBS (Abcam, #ab47003) for 1 hour at 37°C, then washed in 1X PBS-T. Next, slides were incubated with 1:200 anti-WGA-Alexa Fluor 555 and 1:400 anti-rabbit Alexa Fluor 633 (Molecular Probes, #A21070) in 0.05% Tween-20, 1X PBS for 30 minutes at 37°C, and then washed in 1X PBS-T. Sections were incubated for 10 minutes at room temperature with 1:1000 DAPI and then washed in PBS. To quench myocardial autofluorescence, slides were stained with 0.1% Sudan black B in 70% ethanol for 10 minutes followed by rinsing in 1X PBS. Slides were mounted using VectaShield without DAPI (Vector Labs, Burlingame, CA).

Microscopy was performed using a Zeiss LSM510 confocal microscope. To balance a maximal number of cells counted versus identifying EdU+ cells as myocyte vs. non-myocyte, EdU-labeled cells were counted at 20x magnification from 12 fields/section at 7 different levels from apex to base per animal (n=4 mice/group); >20,000 nuclei counted per animal. Fields were chosen at random from different regions of the ventricle on each section to minimize selection bias prior to fluorescence imaging. DAPI-labeled cells were counted at 20x magnification using Image J. Cardiomyocyte nuclei were identified by co-staining with cTnI and were counted separately. The percent of proliferating non-myocyte cells was calculated as the number of EdU-labeled non-myocytes divided by total DAPI-stained non-myocyte nuclei.

Finally, to identify the type of cell labeled with EdU, we co-stained with the Click-iT

EdU Alexa Fluor 488 Imaging Kit and one of the following 3 primary antibodies: (1) 1:500 anti-vimentin (Abcam #ab92547); (2) 1:800 anti-CD31 (Abcam #ab124432); or (3) 1:1000 anti-CD45 (Abcam #ab10558). Primary antibodies were diluted in 3% BSA-1X PBS and were incubated for 1hour at 37°C then overnight at 4°C. The secondary antibody for each was 1:200 anti-rabbit AlexaFluor 633, combined with 1:200 anti-WGA in 0.05% Tween-20 1X PBS. Secondary antibody was incubated for 45 minutes at 37°C. Microscopy was performed using a Zeiss LSM800 Airyscan confocal microscope. Cells were counted from 12 sections/level from 7 different levels of the LV from apex to base from n=4 mice for each antibody. Given that 99.22% of EdU-labeled cells were non-myocytes, all EdU+ cells were considered to be non-myocytes for this analysis. Co-labeling was identified by co-staining of cells for both EdU and the marker of interest.

*Pooled Cardiac Cell Isolation and Immunocytochemistry:* Hearts were exposed by midline thoracotomy, excised and placed in cold perfusion solution (NaCl 113mM, KCl 4.7mM, $Na_2HPO_4$ 0.6mM, $KH_2PO_4$ 0.6mM, $MgSO_4$ 1.2mM, Phenol Red 0.032mM, $NaHCO_3$ 12mM, $KHCO_3$ 10mM, Hepes 10mM, Taurine 30mM, Glucose 10mM, 2,3-Butanedione Monoxime 10mM). Retrograde coronary perfusion was established via aortic cannulation using the Langendorf hanging heart method.(3) The heart was perfused with enzyme buffer (0.48mg/ml collagenase 4, 0.56mg/ml collagenase 2 (Worthington Biochem Corp.) , 0.072mg/ml proteinase XIV (Sigma), in perfusion solution) for 10 minutes. The atria and right ventricle were removed and the LV was minced into small pieces in transfer buffer (perfusion solution + 5mg/mL BSA) and then passed several times through a sterile pipette. The resulting cell suspension was passed through a mesh filter (250 µm) into a 50mL centrifuge tube and incubated for 15 minutes at RT to allow myocytes to pellet by gravity. The pellet was collected as a cell fraction enriched in myocytes. The supernatants from the filtered cell solution was centrifuged at 1500 rpm for 5 minutes at 4°C and then plated in 75mm tissue culture dishes. After 2 hours, dead cells were washed off with 1x PBS and plated cells collected, resulting in a non-myocyte cell population.

To assess if our cell isolation protocol biases towards collection of one type of cardiac non-myocyte, immunocytochemistry was performed. Acid-washed, gelatin coated cover slips were created by washing coverslips in 1M HCl for 6 hours at 60°C with occasional agitation. Coverslips were rinsed with distilled water 4x and then autoclaved. Coverslips were placed individually into wells of a 6-well plate; 2ml of 0.2% gelatin solution was added and incubated for 30 minutes. The gelatin solution was aspirated and 1mL of DMEM F12 +10% FBS, 1%pen/strep was added. Cells were seeded and cultured for 18 hours then fixed in 2mL 4% paraformaldehyde (in 1X PBS) for 10min. Cells were rinsed 3x5 minutes with 1X PBS. Permeabilization was performed with 0.05% Triton X-100 in 1X PBS for 5 minutes at room temperature. Fixative was quenched with 50mM $NH_4Cl$ in 1X PBS for 10 minutes then rinsed 1x5 minutes in 1X PBS. Fixed cells were blocked in 3% BSA (in 1X PBS) and incubated overnight at 4°C. Staining was performed with 1:300 α-vimentin (Abcam, #ab24525), 1:50 α-CD31 (Abcam, #ab9498) and 1:50 α-CD45 (Abcam, #ab10558). Coverslips were washed 3x5 minutes in 1X PBS then once with 3% BSA in 1X PBS. Secondary antibodies were applied (1:400 AlexaFluor-488 goat α-rat IgG (Abcam, #ab150165), 1:400 AlexaFluor-555 goat α-chicken IgG (Abcam, #ab21437), and 1:400 AlexaFluor-647 goat α-rabbit IgG (Abcam, #ab1500083)) and incubated 1 hour at room temperature in a dark chamber. Coverslips were washed 3x5 minutes in 1X PBS then incubated with 1:500 DAPI in water for 10 minutes at room temperature. Coverslips were washed 3x5 minutes in water and then mounted with Vectashield. Confocal microscopy was performed (**Supplementary Figure 2**).

*Protein Isolation and Western Blotting:* Total protein was extracted from flash frozen LV tissue from ~20-week old $PLN^{R9C}$ or age-matched WT mice using RIPA buffer (Millipore). Protein lysates were heated to 95°C, run on 4-12% bis-tris Plus pre-cast gels and transferred to PVDF membranes via wet transfer. For all western blotting, imaging was performed on a Kodak Image Station 4000MM Pro. Total protein was stained using Ponceau S and quantified using Image J. Band densitometry was performed using Image J.

Western blotting for BET family proteins was performed using (1) 1:250 anti-Brd4 (Abcam, #ab128874) with 1:1000 anti-rabbit IgG-HRP (Cell Signaling Technology, #7074); (2) 1:1000 anti-Brd3 (Abcam, #ab71815) with 1:2000 anti-rabbit IgG-HRP; (3) 1:250 anti Brd2 (Cell Signaling, #5848S) with 1:1000 anti-rabbit IgG-HRP; and 1:20,000 anti-vinculin (Abcam, #ab129002) with 1:2000 anti-rabbit IgG-HRP. Primary antibodies were diluted in 5% nonfat milk/1X TBS-T and incubated overnight at 4°C, while secondary antibodies were diluted in 5% nonfat milk/1X TBS-T and incubated for 30 to 60-minutes at 37°C.

Western blotting for NFκB family members was performed using (1) 1:1000 anti-NFκB p65 (Abcam, #ab16502) with 1:1000 anti-rabbit IgG (Cell Signaling Technology, #7074); and (2) 1:1000 anti-NFκB p65 acetyl K310 (Abcam, #ab19870).

*RNA Isolation and RNA-seq:* RNA was extracted from excised LV tissue or pooled non-myocyte and cardiomyocyte fractions using TRIzol (Thermo-Fisher). Quality and quantity was assessed using a TapeStation 2200 (Agilent). Prior to cDNA synthesis, 2 rounds of poly-A selection were performed using oligo dT Dynabeads (Thermo-Fisher). cDNA libraries were constructed from individual mouse samples and RNA-seq libraries were constructed using the Nextera XT DNA

Library Preparation Kit (Illumina). Libraries were sequenced on the Illumina platform, then aligned to the mouse reference sequence mm10 using the STAR aligner software.(4) The total number of reads of each transcript was normalized to one million, with p-values of gene fold-changes determined as previously described.(5)

*Bioinformatics Pipeline for Pathway Analysis:* To confirm the consistency of gene expression changes between mice in different treatment groups, principle components analysis (PCA) was performed on RNA-seq data using the R packages "psych" and "ggplot2" via a custom pipeline. The most significant principle components for each comparison (preDCM and DCM age groups) were determined from Scree plots by a combination of (1) Kaiser-Harris criteria, in which only those principle components with eigenvalues >1 are retained, (2) a Cattell-Scree test, in which only those principle components above the "elbow" (first major inflection point on a Scree plot) are retained, and (3) eigenvalues based on real data that are above a simulated eigenvalue generated from a random data matrix of the same size as our input RNA-seq dataset.

Ingenuity Pathway Analysis (IPA) uses proprietary algorithms and a proprietary database of gene information to generate different analyses based on a set of user-identified genes (http://www.ingenuity.com/products/ipa). We used IPA to identify enriched canonical pathways and to predict possible upstream regulatory molecules from our RNA-seq data. Only genes identified as differentially expressed were uploaded into IPA. We used the following IPA Core Analysis Settings: General Settings, reference set = Ingenuity Knowledge Base (Genes only), relationships to consider = direct and indirect relationships; Data Sources = all; Confidence = experimentally observed only; Species = all. Some gene symbols in our mm10 database were not identifiable by IPA and were consequently not used in the analysis (range 11-14%).

First, canonical pathways were reviewed to assess known biological pathways that were enriched among differentially expressed genes. Specific canonical pathways of interest were assessed for significance based on the nominal p-value as determined by IPA software (right-tailed Fisher exact test). This nominal p-value is a measure of the likelihood that the association between a particular canonical pathway and our differentially expressed genes is due to chance. We considered a nominal p-value of $<0.05$ significant when considering specific pathways of interest. To control for the rate of false discoveries when analyzing all significantly enriched pathways for a given set of genes, statistical significance was based on the p-value corrected for multiple hypothesis testing (Benjamini-Hochberg). In this case, the p-value represents the false-discovery rate (FDR) and are listed as such where appropriate. A FDR $<0.01$ was defined as significant.

IPA also attempts to predict whether a pathway is activated or inhibited with the use of a z-score. The z-score is a standard deviation and represents the confidence with which a pathway can be assigned directionality. In our analyses, we considered a canonical pathway z-score $\geq 2$ (activated) or $\leq -2$ (inhibited) as significant in the setting of a nominal p-value$<0.05$. For many pathways, insufficient data is available to predict activation, and for these pathways, no z-score is given (listed as “uncertain” in our supplementary tables).

In addition to analyzing canonical pathways, we also used the IPA upstream regulator tool.(6) This bioinformatics analysis software attempts to predict the functional status of molecules (called “upstream regulators”) including but not limited to transcription factors, kinases, transmembrane receptors, cytokines and growth factors, based on the known association between these upstream regulators and their downstream targets (the input set of differentially expressed genes). We defined 2 metrics of significance for these analyses: (1) overlap p-value, and (2) z-score. The overlap p-value represents the statistical significance in the overlap between the differentially expressed genes (the known downstream targets) and all genes associated with a particular upstream regulator in the Ingenuity database. We used a p-value threshold of $<0.01$ as significant. However, the p-value of overlap is direction-independent; i.e., the statistical association does not account for whether a gene is up- or down-regulated, which is expected to have substantial impact on the biological relevance of a particular finding. The z-score does account for the direction of activation of the differentially expressed genes in relation to the published literature with respect to a particular upstream regulator, thereby allowing a prediction of whether the upstream regulator is activated or inhibited. Four possibilities exist: (1) a gene known to be activated by an upstream regulator that is up-regulated in the dataset predicts activation of the upstream regulator; (2) a gene known to be activated by an upstream regulator that is down-regulated in the dataset predicts inhibition of the upstream regulator; (3) a gene known to be inhibited by an upstream regulator that is up-regulated in the dataset predicts inhibition of the upstream regulator; (4) a gene known to be inhibited by an upstream regulator that is down-regulated in the dataset predicts activation of the upstream regulator. The z-score accounts for both the total number of genes differentially expressed and the direction of activation; a z-score $\geq 2$ (activated) or $\leq -2$ (inhibited) was considered significant.

For more details on calculation of p-values by IPA and the IPA upstream analysis tool, the interested reader is referred to the following online primer materials:

1. “Understanding the p-value of overlap statistic in IPA” tutorial video: (https://tv.qiagenbioinformatics.com/video/19605716/understanding-the-p-value-of).

2. The “Ingenuity upstream regulator analysis in IPA” White paper: (http://pages.ingenuity.com/rs/ingenuity/images/0812

upstream_regulator_analysis_whitepaper.pdf)

3. Multiple videos on the tutorial page supporting the upstream analysis tool:

(https://tv.qiagenbioinformatics.com/channel/13382329/tutorials).

### SUPPLEMENTARY TABLES

**Supplementary Table 1:** see attached excel file. Differentially expressed (D.E.) genes in preDCM aged mice treated with JQ1 or vehicle from 6 to 8-weeks of age were compared as follows: WT-JQ1 vs. WT-vehicle (sheet 1), PLN-vehicle vs. WT-vehicle (sheet 2), PLN-JQ1 vs. WT-vehicle (sheet 3) and PLN-JQ1 vs. PLN-vehicle (sheet 4). Each sheet displays gene symbol and gene name/description, normalized RNA-seq read counts for the D.E. genes within the indicated groups, and fold-change plus the associated p-values for each comparison between mice within groups. We have listed the normalized read counts for the D.E. genes within each group for ALL mice in the study so that the interested reader may assess what happened to D.E. genes in different groups of mice. The normalized reads used for calculating fold change and p-values are highlighted within each table. Mouse nomenclature is as follows: “1WTjq” refers to WT JQ1-treated mouse #1, and so forth.

**Supplementary Table 2:** see attached excel file. Differentially expressed genes expressed in the LV of preDCM (8-week old) mice were subjected to IPA as described in the Supplementary Methods. Significantly enriched ingenuity canonical pathways (defined as FDR <0.01) are shown for the comparison of WT-JQ1 vs. WT-vehicle (sheet 1), PLN-vehicle vs. WT-vehicle (sheet 2), PLN-JQ1 vs. WT-vehicle (sheet 3) and PLN-JQ1 vs. PLN-vehicle (sheet 4). The p-value represents the nominal p-value for a given pathway, while the FDR is the Benjamini-Hochberg p-value corrected for multiple hypothesis testing (see Supplementary Methods). Differentially expressed pathway genes are the percent of genes within a pathway that are differentially expressed in the comparison. The z-score represents the directionality of the pathway (positive = activated pathway; negative = inhibited pathway; uncertain = gene expression data are insufficient to predict directionality; see Supplementary Methods).

**Supplementary Table 3:** see attached excel file. Differentially expressed genes expressed in the LV of preDCM (8-week old) mice were subjected to the IPA Upstream Regulator analysis tool (described in the Supplementary Methods). Significantly enriched regulators (defined as $P<0.01$ AND $Z\pm2$) are shown for the comparison of WT-JQ1 vs. WT-vehicle (sheet 1), PLN-vehicle vs. WT-vehicle (sheet 2), PLN-JQ1 vs. WT-vehicle (sheet 3) and PLN-JQ1 vs. PLN-vehicle (sheet 4). Regulators are separated by type and differentially expressed genes that drive the result are listed ("target molecules in dataset").

**Supplementary Table 4:** see attached excel file. Differentially expressed (D.E.) genes in mice with overt DCM (20-weeks of age) after 12 weeks of treatment with JQ1 or vehicle were compared as follows: WT-JQ1 vs. WT-vehicle (sheet 1), PLN-vehicle vs. WT-vehicle (sheet 2), PLN-JQ1 vs. WT-vehicle (sheet 3) and PLN-JQ1 vs. PLN-vehicle (sheet 4). Each sheet displays gene symbol and gene name/description, normalized RNA-seq read counts for the D.E. genes within the indicated groups, and fold-change plus the associated p-values for each comparison between mice within groups. We have listed the normalized read counts for the D.E. genes within each group for ALL mice in the study so that the interested reader may assess what happened to D.E. genes in different groups of mice. The normalized reads used for calculating fold change and p-values are highlighted within each table. Mouse nomenclature is as follows: "1WTjq" refers to WT JQ1-treated mouse #1, and so forth.

**Supplementary Table 5:** see attached excel file. Differentially expressed genes expressed in the

LV of DCM (20-week old) mice were subjected to IPA as described in the Supplementary Methods. Significantly enriched ingenuity canonical pathways (defined as FDR <0.01) are shown for the comparison of WT-JQ1 vs. WT-vehicle (sheet 1), PLN-vehicle vs. WT-vehicle (sheet 2), PLN-JQ1 vs. WT-vehicle (sheet 3) and PLN-JQ1 vs. PLN-vehicle (sheet 4). The p-value represents the nominal p-value for a given pathway, while the FDR is the Benjamini-Hochberg p-value corrected for multiple hypothesis testing (see Supplementary Methods). Differentially expressed pathway genes are the percent of genes within a pathway that are differentially expressed in the comparison. The z-score represents the directionality of the pathway (positive = activated pathway; negative = inhibited pathway; uncertain = gene expression data are insufficient to predict directionality; see Supplementary Methods).

**Supplementary Table 6:** see attached excel file. Differentially expressed genes expressed in the LV of preDCM (8-week old) mice were subjected to the IPA Upstream Regulator analysis tool (described in the Supplementary Methods). Significantly enriched regulators (defined as P<0.01 AND Z±2) are shown for the comparison of WT-JQ1 vs. WT-vehicle (sheet 1), PLN-vehicle vs. WT-vehicle (sheet 2), PLN-JQ1 vs. WT-vehicle (sheet 3) and PLN-JQ1 vs. PLN-vehicle (sheet 4). Regulators are separated by type and differentially expressed genes that drive the result are listed (“target molecules in dataset”).

**SUPPLEMENTARY FIGURES**

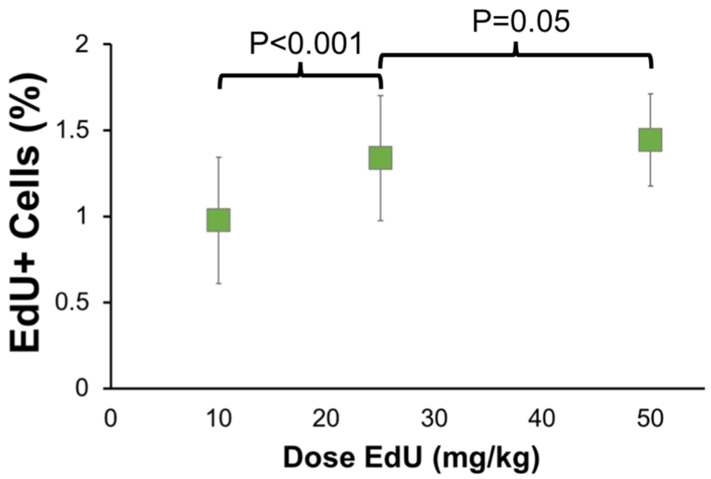


**Supplementary Figure 1. EdU dose titration in the heart.** The optimal dose of EdU for labeling cardiac cells is unknown. EdU was injected to n=4 wild type mice on consecutive days prior to sacrifice with the doses indicated. Hearts were excised, fixed, paraffin embedded and stained using the Click-iT EdU labeling kit as described in the methods section. Slides containing 6 sections each from 7 different levels evenly distributed from apex to base were stained. Cells from 6 randomly selected 10X magnification regions of each slice (36 regions of myocardium per slide) were counted and EdU+ cells quantified. An average of 77,334±5771 cells were counted per mouse.

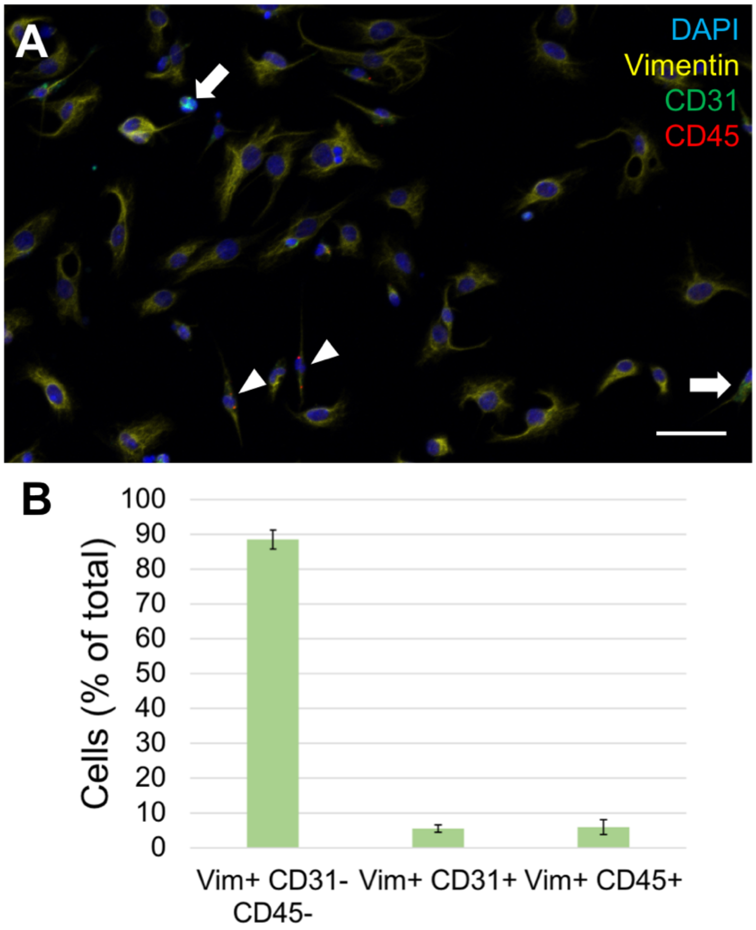


**Supplementary Figure 2. Immunocytochemistry of fixed, pooled cardiac non-myocytes. (A)** Sample confocal image of plated cardiac non-myocytes stained with DAPI, α-vimentin, α-CD31 (white arrows) and α-CD45 (white arrowheads). Nearly all non-myocytes were vimentin-positive. A small portion were also CD31- or CD45-positive as well. **(B)** Quantification of cell counts from ~1200 nuclei from n=5 mice showing that nearly 90% of non-myocyte cells were vimentin+/CD31-/CD45-, thus demonstrating that our cardiac non-myocyte population is heavily enriched for cardiac fibroblasts.

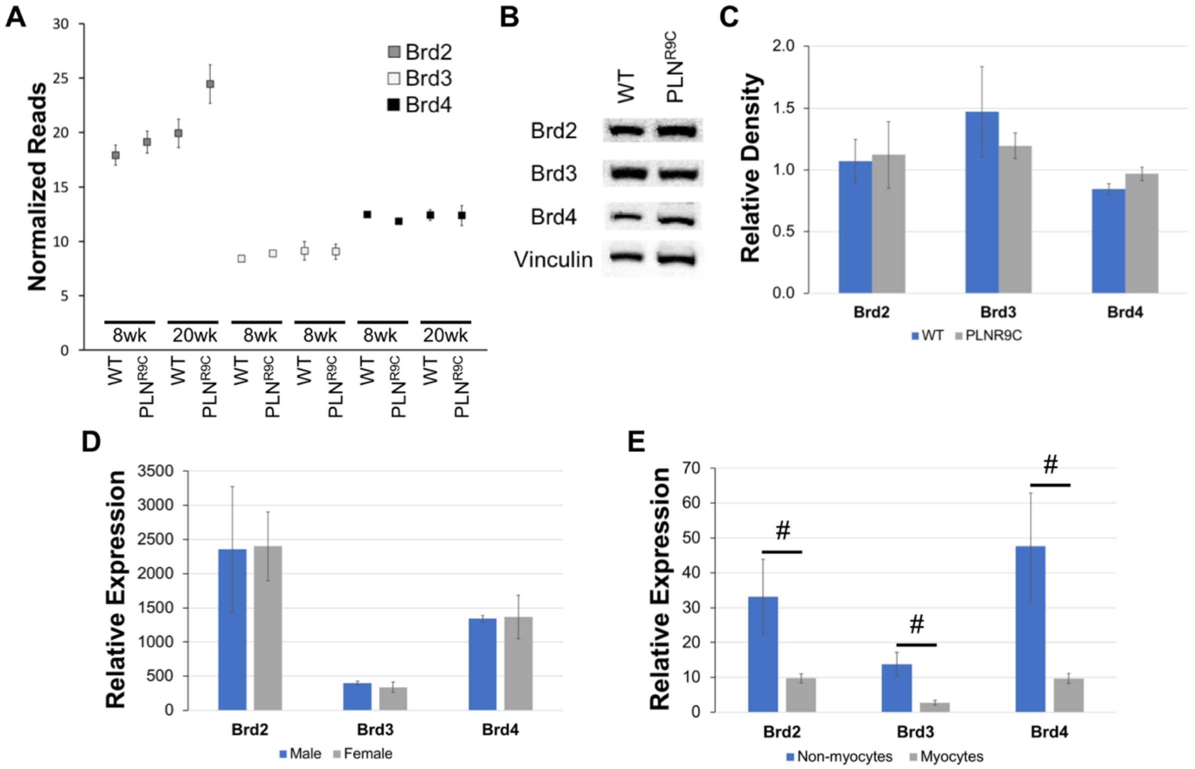


**Supplementary Figure 3. BET genes are expressed in PLN$^{R9C}$ and wild type (WT) mice. (A)** BET bromodomain genes were expressed at all ages and stages of disease. Data are expressed as the average normalized read count of n=3 mice ± SD from RNA-seq libraries. Data were confirmed by Q-PCR (not shown). *Brdt* was not expressed in the heart (data not shown). **(B)** Representative western blots and **(C)** densitometry (n=3) showed robust BET expression in LV tissue lysates. **(D)** Cardiac BET gene expression was comparable between males and females (droplet digital PCR on whole LV tissue samples relative to 18s rRNA x $10^{-6}$; n=3)**. (E)** BETs were expressed in both cardiac non-myocytes as well as cardiomyocytes in WT and PLN$^{R9C}$ mice at 18-weeks of age, with predominant expression of all 3 genes in the cardiac non-myocyte population (digital droplet PCR on isolated cardiac cell fractions normalized to 18s rRNA x $10^{-6}$; n=6; #p<0.001).

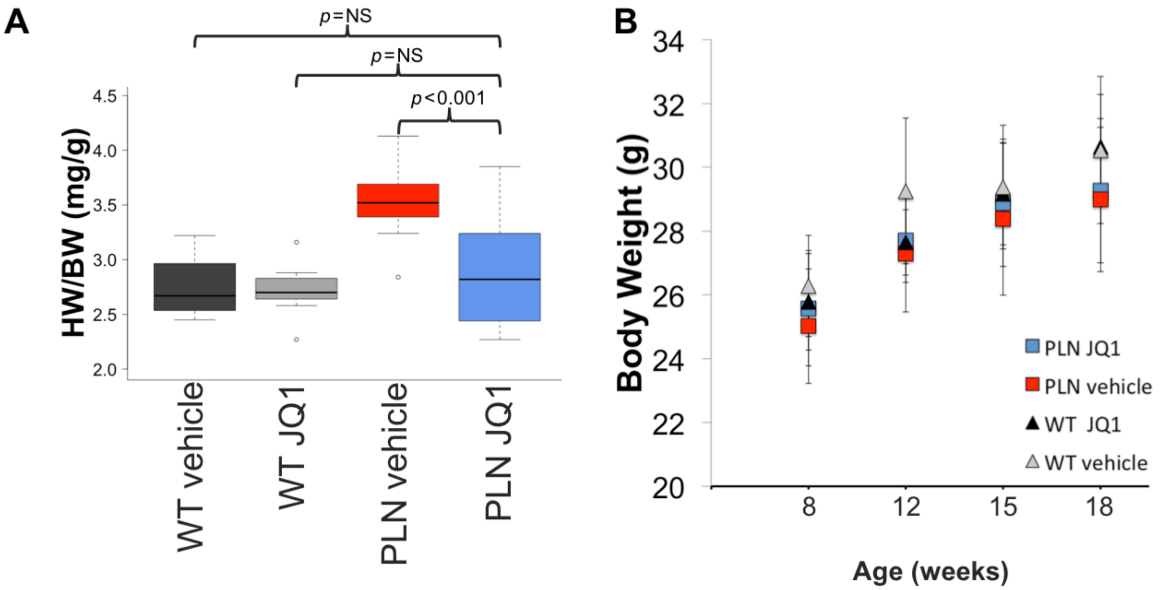


**Supplementary Figure 4. BET inhibition prevents LV hypertrophy. (A)** Heart weight (HW), as estimated by echocardiography, was normalized to body weight (BW) prior to sacrifice, and showed that JQ1 treatment in $PLN^{R9C}$ mice prevented LV hypertrophy. JQ1 did not affect cardiac hypertrophy in wild type mice. **(B)** BW increased with time ($P<0.001$) consistent with normal growth in all age groups. There was a non-significant trend to lower body weight in $PLN^{R9C}$ mice at 18-weeks of age. JQ1 had no effect on body weight.

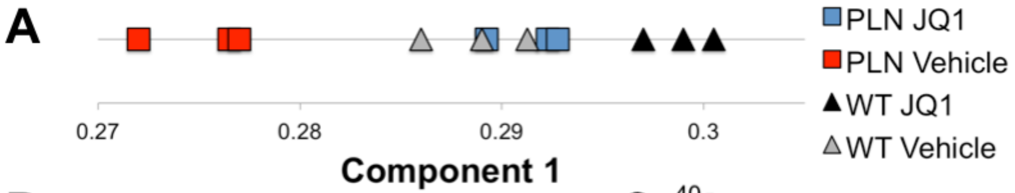


**B**

| | Component 1 | Component 2 |
|---|---|---|
| Standard deviation | 6.3 | 0.7 |
| Proportion of variance | 97.3% | 1.2% |
| Cumulative variance | 97.3% | 98.5% |

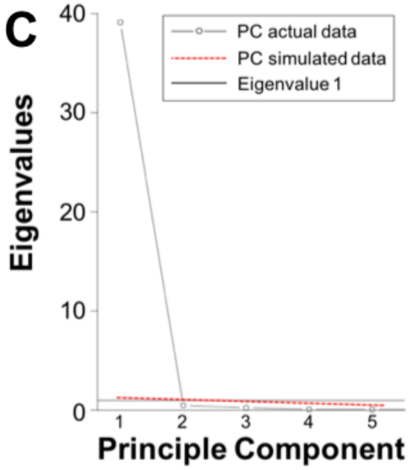


**Supplementary Figure 5. Principle Components Analysis (PCA) in preDCM mice.** PCA was performed on whole LV RNA-seq data to assess global gene expression patterns in response to JQ1 or vehicle treatment. **(A)** Mice within each treatment arm grouped very closely together by gene expression with $PLN^{R9C}$ vehicle-treated mice demonstrating the most globally divergent gene expression program. $PLN^{R9C}$ JQ1-treated mice have a gene expression pattern that overlapped WT vehicle-treated mice. **(B)** PCA revealed that component 1 constituted >97% of the variance observed across the 4 groups of mice, thus PCA data are plotted in 1-dimension in panel A. **(C)** PCA scree plot and parallel analysis of gene expression demonstrates that component 1 is the only component that (1) meets Kaiser-Harris criteria (eigenvalue >1), (2) is above the inflection point (the Cattell Scree test), and (3) is greater than the simulated eigenvalue using random data, thus supporting that component 1 accounts for nearly all of the variance observed between 8-week old mice irrespective of treatment.

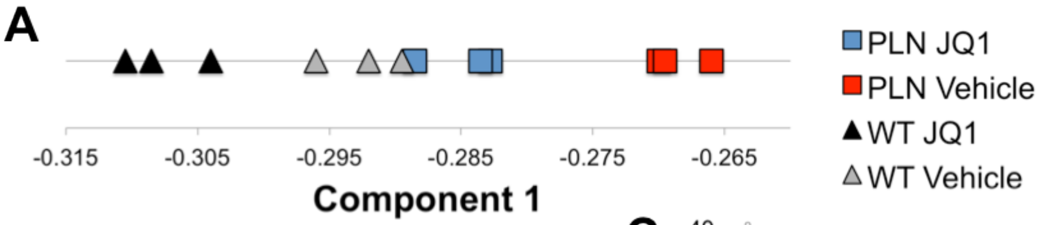


B

| | Component 1 | Component 2 |
|---|---|---|
| Standard deviation | 6.4 | 1.1 |
| Proportion of variance | 94.9% | 2.8% |
| Cumulative variance | 94.9% | 97.7% |

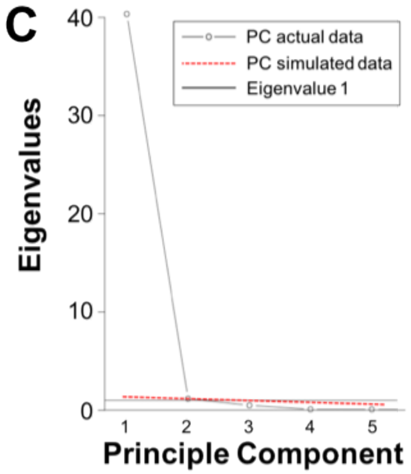


**Supplementary Figure 6. Principle Components Analysis (PCA) in mice with advanced DCM.** PCA was performed on whole LV RNA-seq data to assess global gene expression patterns in response to JQ1 or vehicle treatment. **(A)** Mice within each treatment arm were grouped closely by gene expression with $PLN^{R9C}$ vehicle-treated mice demonstrating the most globally divergent gene expression program. Gene expression in $PLN^{R9C}$ JQ1-treated mice was most similar to WT vehicle-treated mice. **(B)** PCA revealed that component 1 constituted ~95% of the variance observed across the 4 groups of mice, thus PCA data are plotted in 1-dimension in panel A. **(C)** PCA scree plot and parallel analysis of gene expression demonstrates that component 1 is the only component that (1) meets Kaiser-Harris criteria (eigenvalue >1), (2) is above the inflection point (the Cattell Scree test), and (3) is greater than the simulated eigenvalue using random data, thus supporting that component 1 accounts for nearly all of the variance observed between 20-week old mice irrespective of treatment.

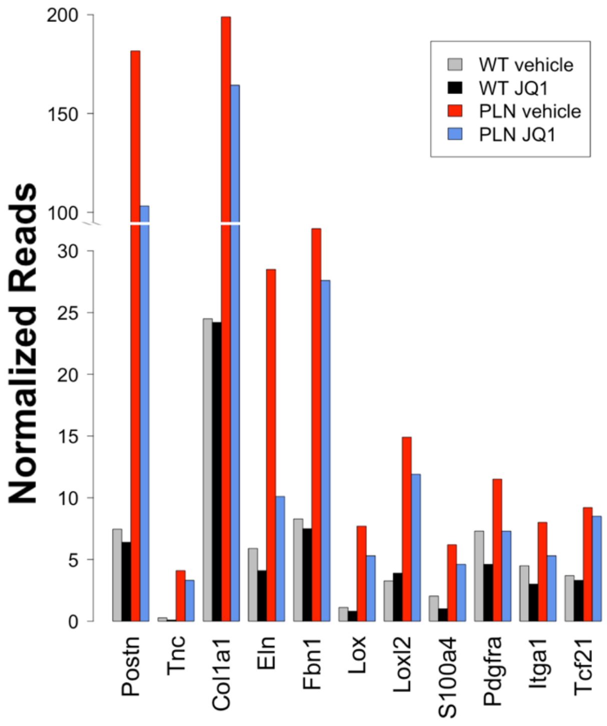


**Supplementary Figure 7. Myofibroblast gene activation in PLN$^{R9C}$ hearts.** RNA-seq normalized read counts for the genes shown from whole LV heart tissue samples. These genes are characteristic of myofibroblasts, and are drastically upregulated in PLN$^{R9C}$ hearts with DCM. There was uniform, albeit incomplete, suppression of myofibroblast gene expression in PLN$^{R9C}$ mice treated with JQ1 ($p<0.001$ for all comparisons of PLN$^{R9C}$ vehicle vs. WT vehicle, and for all comparisons of PLN$^{R9C}$ JQ1 vs PLN$^{R9C}$ vehicle except Tcf21). Tcf21 is a fibroblast-specific transcription factor expressed in all types of cardiac fibroblasts and is shown here to confirm the presence of fibroblasts in the sample; the increased level of Tcf21 in PLN$^{R9C}$ is believed to be due to the increased absolute number of fibroblasts in these hearts.

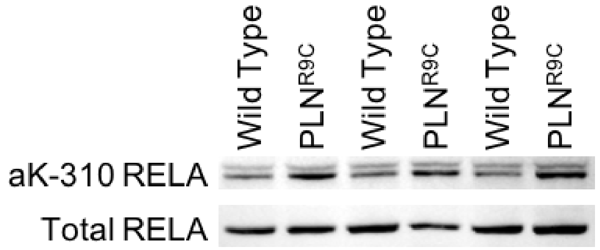


**Supplementary Figure 8. NFκB is activated in PLN$^{R9C}$ hearts.** Western blots showing total and acetyl lysine 310 (aK310) RELA levels from whole LV tissue lysates in 22-week old mice with DCM relative to age-matched wild type mice demonstrating increased NFκB activation as suggested by higher aK310 levels.Files
